# Quantification of the effects of single nucleotide variants in NKX2.1 transcription factor binding sites

**DOI:** 10.64898/2026.02.27.708450

**Authors:** Freyda Lenihan-Geels, Sebastian Proft, Martin Bommer, Udo Heinemann, Dominik Seelow, Robert Opitz, Heiko Krude, Markus Schuelke, Maria Małecka

## Abstract

Transcription factors recognise and bind specific DNA sequence patterns in promoters and enhancers thereby regulating gene expression. Variations in the DNA sequence of transcription factor binding sites (TFBSs) can alter gene regulation and may disrupt development. The transcription factor NKX2.1 is a crucial regulator of thyroid, lung, and neural development. Mutations in its coding gene *NKX2-1* may cause choreoathetosis and congenital hypothyroidism with or without pulmonary dysfunction (CAHTP, OMIM #610978). Most genetically solved patients carry mutations in the coding regions of *NKX2-1* that affect DNA binding, while the majority of patients with CAHTP-like symptoms do not carry mutations in the *NKX2-1* coding sequence. We hypothesise that variations in the DNA-sequence at promoter or enhancer sites to which the transcription factor NKX2.1 binds could cause disease as well. We employed EMSA-seq to quantify the effects of genetic variation on NKX2.1 binding strength and used this data to train neural network models to forecast the influence of DNA variation on NKX2.1 binding. We validated our models using microscale thermophoresis, X-ray crystallography, and publicly available ChIP-seq data sets. The neural networks were able to detect TFBSs in ChIP-seq data and can thus be used to evaluate whole genome sequencing data of CAHTP-patients in order to prioritise potential disease-causing variants in regulatory elements.

**Graphical abstract:** 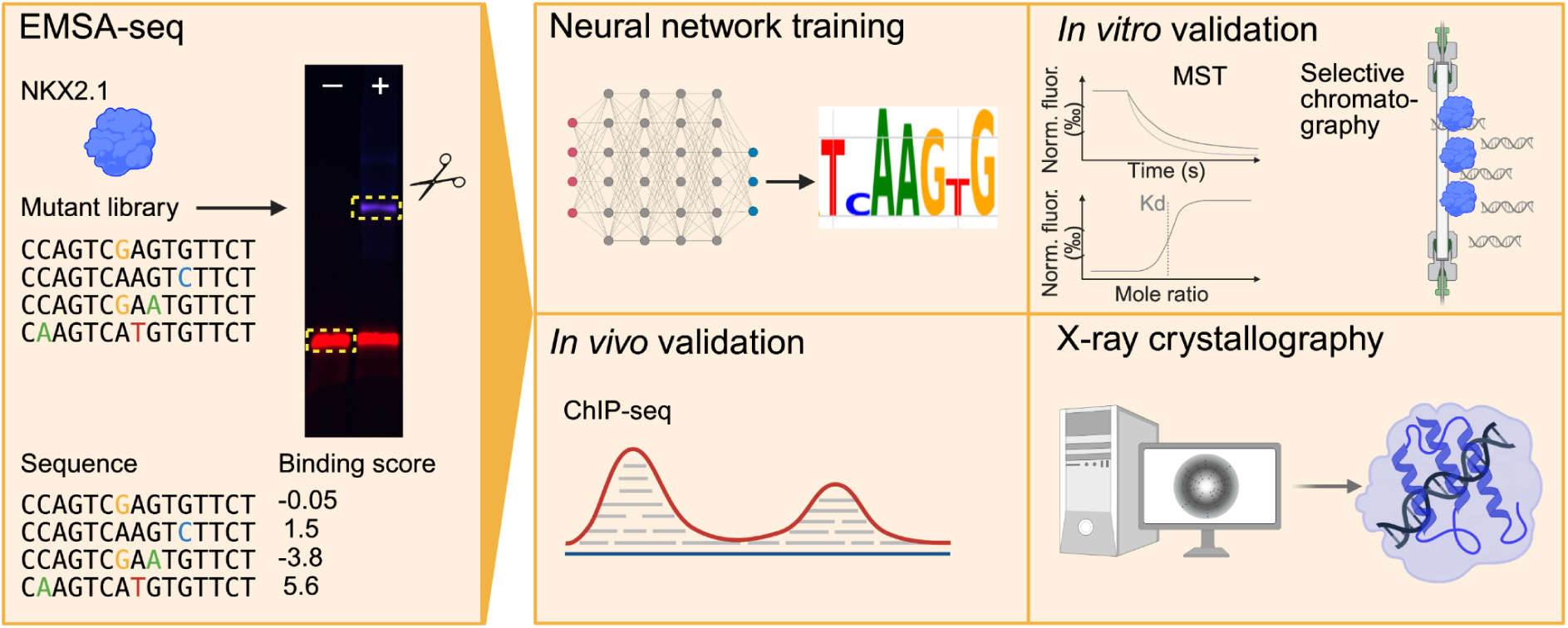

## Introduction

Transcription factors (TFs) are key regulators of cell maintenance and differentiation. They bind to specific sites in promoters, enhancers, or silencers thereby controlling and fine-tuning gene expression [1]. Understanding the DNA-binding preferences of TFs for their transcription factor binding sites (TFBSs), and the effects of single nucleotide variants (SNVs) on their binding strength will provide insight into gene regulation and dysregulation in health and disease. Whole exome sequencing studies in classic Mendelian disorders detect causative disease mutations only on an average of 32% [2]. Given this, we hypothesise that the remaining mutations must reside in the regulatory genome, for example at TFBSs in the promoter or enhancer regions of genes. Due to the complexity and diversity of the human TFs, each TF must be studied individually to understand its binding preferences and dynamics and to predict potential effects of variants in their DNA binding sites.

Advances in sequencing technology and its decreasing cost have made whole genome analysis a viable diagnostic tool. However, the reliable identification of larger numbers of regulatory, disease-causing variants in human TFBSs is still lacking. Rare examples for such variants include point mutations associated with pancreatic agenesis and polyaxial polydactyly. Specifically, SNVs in the *PTF1A* enhancer were found to disrupt binding of two TFs, FOXA2 and PDX1, which are essential for pancreatic development [3]. Point mutations associated with polyaxial polydactyly were found to affect the binding of TFs from the E26 transformation-specific (ETS) family in the zone of polarising activation regulatory sequence that controls sonic hedgehog (*SHH*) expression during limb development [4,5]. Variants in ETS binding sites that increase TF:DNA binding affinity by just 9% relative to the wildtype sequence, are sufficient for ectopic *SHH* expression and the formation of extra digits [5]. The investigation of familial disorders for which no coding variants can be found that would explain the phenotype will require the screening of thousands of regulatory variants and prioritising a subset of candidate variants based on current knowledge of regulatory regions that might contribute to the observed phenotypes. Such a prioritisation requires pedigree construction via whole genome sequencing of multiple affected and unaffected family members and prior knowledge of the exact genomic position of promoters and enhancers. Therefore, prior to extensive experimental testing, we need accurate models of TF binding for the prioritisation of potential candidate SNVs that may cause altered TF binding.

The presently most commonly employed TF binding models are position weight matrices (PWMs), which determine the probability of a nucleotide occurring at each position of the binding site [6]. However, PWMs are not intended for predicting altered binding due to variations in the DNA. PWMs assume that each nucleotide contributes to binding independently, and only transcription factor flexible models, consider relationships between neighbouring dinucleotides [7]. Recently, deep learning methods have been applied to the task of modelling TF binding [8–10]. Deep learning models, especially neural networks, are able to learn complex relationships between DNA sequence and TF binding, taking also into account interdependencies even between distal nucleotides. These models contain many layers and the extraction of meaning from these networks for the inference of biological mechanisms is difficult. Tools such as *Deep Learning Important FeaTures* (DeepLIFT) [11] and *Self-attention for HierArchical Feature Transfer and Selection* (SHAFTS) [12] can be used to decipher attributions and delineate the importance of different nucleotides and positions at a predicted TFBSs.

*In vitro* experimental methods allow the analysis of TF:DNA interactions without the interference of other levels of transcriptional regulation, such as chromatin accessibility and chemical DNA modifications (e.g. CpG island methylation). Methods such as microscale thermophoresis (MST) [13,14] and isothermal calorimetry (ITC) [15] can be used to measure the physicochemical affinity between TFs and DNA at TF binding sites. These methods can only investigate the binding of a single DNA sequence at a time. Conversely, *in vitro* methods such as *ElectroMobility Shift Assay* sequencing (EMSA-seq) [16–18], *High Throughput Systemic Evolution of Ligands by EXponential enrichment* (HT-SELEX) [19,20], *Protein Binding Microarrays* (PBMs) [21–24], and *Mechanically Induced Trapping Of Molecular Interactions* (MITOMI) [25–27] can determine relative TF binding strengths to many DNA sequences in parallel. While these methods do not provide a direct measurement of TF:DNA binding energies, they can be used to compare the relative binding strength of different DNA sequences, and in turn model TF binding specificities.

In a recent study of n = 101 patients with the complete or partial characteristic disease phenotype associated with *NKX2-1* mutations (e.g. choreoathetosis and congenital hypothyroidism with or without pulmonary dysfunction, CAHTP, OMIM #610978) [28–49], 73% had remained unsolved [48]. The NKX2.1 protein is a homeodomain TF that plays an important role in development and maintenance of thyroid, lung, and brain tissues, especially the basal ganglia, e.g. mainly the *Pallidum* [50,51]. NKX2.1, like other homeodomain TFs, recognises 5-7 bp sites in the DNA and binds *via* its DNA-binding domain (DBD) of its homeodomain [52]. Of the *NKX2-1* variants that cause CAHTP, several affect the homeodomain and thus impede the DNA binding strength of the protein [28,32,33,42,47,48]. Inversely, it is plausible that pathogenic variants at NKX2.1 binding sites on the DNA may also disrupt regulatory capabilities and cause disease. Indeed, variants in NKX2.1 binding sites have been associated with chronic obstructive pulmonary disease [53,54] and Hirschsprung’s disease [55].

PWM models deduced from ChIP-seq data of mouse lung, lung adenocarcinoma, and basal ganglia cells [56] have been used to describe the NKX2.1 binding site, and these models have been validated in human lung cancer cell lines [57]. However, there is no publicly available *in vitro* binding data for this TF. Hence, we set out to quantify the effects of SNVs and of combinations thereof in the NKX2.1 binding site to create TFBS models that would predict the effect of DNA sequence variation on NKX2.1 binding affinity. We employed EMSA-seq to study the effects of one or more SNVs in the TFBS on DNA binding strength in high throughput. We then trained convolution neural networks (CNNs) on the EMSA-seq data to predict the enrichment of DNA sequences upon NKX2.1 binding. Using `tangermeme` [58], we assigned attributions and importance for protein binding to different nucleotides in the DNA sequence. We compared and contrasted the quantitative EMSA-seq findings with binding affinity measurements from MST, as well as X-ray crystallography structural data and `AlphaFold` structure predictions of NKX2.1:DNA interactions. The binding model presented here can be used in a clinical setting, to assess the effects of variants in promoters and enhancers suspected of affecting NKX2.1 binding, particularly when no deleterious coding variants are found.

## Materials and methods

### Recombinant protein production

Recombinant protein expression was performed using an *Escherichia coli* expression system for two distinct constructs of *NKX2-1*, differing in length and fusion tags depending on their downstream application. Briefly, the gene encoding the 67 amino acid DNA-binding domain of NKX2.1 (truncated 1, 67 amino acids, aa192-258 of NP_001073136.1) was cloned into the modified pET-28a vector, incorporating an N-terminal TRX-His6-tag followed by a Tobacco Etch Virus (TEV) cleavage site, and a C-terminal green fluorescent protein (GFP) as a fusion protein. Concurrently, a truncated version of the gene encoding the NKX2.1 DNA-binding domain (truncated 2, 59 amino acids, aa192-250 of NP_001073136.1) was cloned into the modified pET-28a vector, incorporating an N-terminal TRX-His6-tag followed by a TEV cleavage site.

Each construct was then separately transformed into *Escherichia coli* BL21 Star™ (DE3, Invitrogen, Carlsbad, CA, USA) chemocompetent cells. Positive transformants for each variant were selected on Luria-Bertani (LB) agar supplemented with 50 µg/ml kanamycin. For protein expression of both variants, kanamycin-supplemented Terrific Broth (TB) was inoculated with 5% of overnight cultures and grown at 37°C with constant shaking until the optical density (OD_600_) reached 0.8. Protein expression was then induced by adding β-D-1-thiogalactopyranoside (IPTG) to a final concentration of 0.5 mM, and the cells were further incubated at 18°C with constant shaking overnight. Bacterial cell pellets were then harvested by centrifugation at 5,000 *x g* for 20 minutes and frozen.

### Protein purification

Bacterial cell pellets were resuspended in H0 buffer [25 mM MES-NaOH ph 6.5, 500 mM NaCl, 40 mM imidazole, 10% glycerol] supplemented with 1 mg/ml lysozyme, 1 ml of DNase I and 1 protease inhibitor cocktail tablet (Roche, Basel Switzerland) per 50 ml of buffer. Cells were lysed by sonication on ice and the lysate was clarified by centrifugation at 20,000 *x g* for 30 minutes at 4°C. The supernatant, containing the soluble NKX2.1 protein, was loaded onto a 5 ml HisTrap^TM^ Crude column (Cytiva, Freiburg, Germany) pre-equilibrated with H0 buffer. The column was washed with 10 column volumes (CV) of P7 buffer [1 M sodium phosphate buffer pH 7.0, 10% glycerol] and 10 CV H0 to remove residual DNA and non-specifically bound proteins. The His-tagged protein was eluted with H500 [25 mM MES-NaOH pH 6.5, 500 mM NaCl, 500 mM imidazole, 10% glycerol]. Fractions containing the target protein were pooled, dialyzed *versus* dialysis buffer [25 mM MES-NaOH ph 6.5, 200 mM NaCl, 10% glycerol], and the His-tag was cleaved by the addition of TEV protease at a 1:100 (w/w) ratio during overnight dialysis against fresh dialysis buffer at 4°C. To remove the cleaved His-tag and protease, the sample was then passed through a 5 ml Uni SP cartridge (BioRad, USA) equilibrated with 80% of Q1 [25 mM MES-NaOH pH 6.5,10% glycerol] and 20% of Q10 [25 mM MES-NaOH pH 6.5, 1 M NaCl, 10% glycerol], the column was washed with 10 CV of equilibration buffer and then the variant NKX2.1 protein was step-eluted at 50% of Q10. Finally, the protein was concentrated using Amicon^TM^ Ultra Centrifugal Filter, 5 kDa MWCO (Millipore, Darmstadt, Germany) and further purified by size exclusion chromatography using a HiLoad 16/600 Superdex® 75 pg (Cytiva) pre-equilibrated with S1 buffer [25 mM MES-NaOH ph 6.5, 500 mM NaCl, 10% glycerol], or S2 buffer [25 mM MES-NaOH ph 6.5, 185 mM NaCl, 10% glycerol], depending on the downstream usage of sample, and fractions corresponding to the monomeric protein, verified by SDS-PAGE electrophoresis, were collected for downstream applications.

For microscale thermophoresis (MST) analysis, the purified protein (truncated 1 with GFP tag) was dialysed after the size-exclusion step and diluted in an S1 buffer [25 mM Hepes pH 7.5, 185 mM NaCl, 1% 1,3-propanediol and 0.05% Tween20] to a concentration of 50 µM. Aliquots of this stock were then snap-frozen in liquid nitrogen and stored at -80°C as single-experiment stocks to maintain protein integrity and prevent photo-bleaching of GFP.

### Preparation of double-stranded DNA and protein sample for MST assay

Oligonucleotides consisting of 24-mer sequences featuring a 4-nucleotide DNA core motif (CAAG) flanked by wildtype 5’ and 3’ sequences [5’-CACTGCCCAGT**CAAG**TGTTCTTGA-3’] (**Supplementary Table S1: oligo 1**), and containing single variants within the motif, were custom-synthesised and purchased from Eurofins Genomics (Ebersberg, Germany). This sequence is a well-characterised NKX2.1 binding site from the rat thyroglobulin promoter [59]. Individual forward and complementary reverse strands were initially diluted in nuclease-free water. Their precise concentrations were determined using a NanoDrop One spectrophotometer (Thermo Scientific, Waltham, MA, USA) at 260 nm. For duplex formation, equimolar amounts (1:1 ratio) of the forward and reverse strands were mixed to achieve a final duplex concentration of 200 nM. The annealing process involved heating the mixture to 60°C for 10 minutes, followed by slow cooling to 4°C at a rate of 1°C per 4 minutes, using a thermocycler. Successful formation and concentration of the resulting DNA duplexes were verified by measuring the absorbance at 260 nm using a NanoDrop One spectrophotometer.

### Microscale thermophoresis

MST was employed to quantify the binding affinity of NKX2.1 to DNA fragments containing variations in the binding motif of the DNA duplices. All MST experiments were performed using Nano Temper Monolith NT.115 (NanoTemper Technologies, Munich, Germany). The NKX2.1 was intrinsically fluorescent *via* a fusion protein (GFP). Immediately before MST measurements, individual NKX2.1-GFP aliquots were thawed on ice and diluted to a constant concentration of 20 nM in cooled MST assay buffer [25 mM Hepes pH 7.5, 185 mM NaCl, 1% 1,3-propanediol, 0.05% Tween 20]. A series of 16-point, 1:1 serial dilutions of each unlabelled DNA duplex variant was prepared in nuclease-free water, starting from 200 nM. For each measurement, equal volumes of the labeled protein and diluted DNA were mixed, resulting in a final concentration of 10 nM for the protein and a corresponding range for the DNA. Samples were loaded into standard treated Monolith® Capillaries and allowed to equilibrate for 2 minutes at 10°C. MST measurements were performed using the blue channel at 20% excitation power, with 3 replicates for each concentration. Data were analysed using the MO.Affinity Analysis Software v2.3 (NanoTemper Technologies). The normalised fluorescence (Fnorm) from the thermophoresis traces was plotted against the logarithm of the varying DNA concentrations, and dissociation constants (K_d_) were determined by fitting the data to a one-site binding model using the law of mass action assuming that the rate of association and dissociation was proportional to the concentration of its reactants.

### Preparation of double stranded DNA and protein sample for crystallisation

Custom-synthesised 12-mer oligonucleotides were purchased from Eurofins Genomics. The sequences were designed with a central four-nucleotide motif (and variations thereof), flanked by conserved 5’ and 3’ regions [5’-AGT**CAAG**TGTTC-3’] (**Supplementary Table S1: oligo 2**). This design allowed the testing of the canonic target motif and of single nucleotide variants therein. To prepare the double-stranded DNA duplexes, complementary forward and reverse strands were first reconstituted in nuclease-free water and then annealed according to the standard protocol described above. Following annealing, NaCl was added to each duplex solution to a final concentration of 100 mM to prevent protein precipitation during complex formation. For the complex assembly, the purified protein obtained from size-exclusion chromatography was dialysed against buffer [25 mM MES pH 6.5, 200 mM NaCl, and 10% glycerol]. The protein:DNA complex was subsequently formed by mixing the protein with the DNA duplex at a 1.2:1 molar ratio. After a 15-minute incubation period, the mixture was concentrated using a 3 kDa molecular weight cut-off (MWCO) centrifugal filter (Millipore). The final concentration of the protein-DNA complex used for crystallisation ranged from 45 to 55 mM.

### Crystallisation

Crystallisation was performed with truncated 2, the shorter NKX2.1 variant without a GFP tag. Screening of the protein:DNA complex was performed using the hanging drop vapor diffusion method at 20°C. Initial trials were set up using sparse matrix screens, such as the JCSG+ Suite and PACT Premier (Molecular Dimensions, Rotherham, UK). Crystallisation drops were prepared by mixing the protein:DNA complex solution with the reservoir solution in a 1:1 or 2:1 ratio (v/v) on a siliconised coverslip, with a typical total drop volume of 2 µl. The coverslip was then inverted and sealed over a reservoir containing 500 µl of the precipitant solution. The sealed plates were incubated and monitored for crystal growth over a period ranging from several days to weeks. After identifying initial hits, conditions were optimised by preparing fine screens that varied the concentrations around 50% PEG 4K, 1 M NaCl, and 100 mM MgCl₂. This optimization was performed at two pH points, using succinate (pH 5.5) and citrate (pH 4.6) buffers, to improve crystal quality for each complex.

### Data collection and processing

For X-ray diffraction analysis, crystals were harvested using nylon loops, briefly soaked in a cryoprotectant solution (reservoir solution supplemented with 20% PEG 4000 and 20% PEG 200), and flash-cooled in liquid nitrogen. A complete data set was collected at 100 K on beamline P13 at the EMBL Hamburg PETRA III storage ring (DESY, Hamburg, Germany). Diffraction data were recorded on a PILATUS 6M detector, with an oscillation angle of 0.1^∘^ per frame. The diffraction images were subsequently processed, indexed, integrated, and scaled using the XDS software package. The structure was solved by molecular replacement (MR), initially using an NMR-derived structure as a search model (PDB ID: 2L9R), and subsequently with a solved crystal structure of the complex. The final structures were refined using the PHENIX suite.

### Model generation with AlphaFold

Computational modelling of the structural complexes between NKX2.1 and the specific DNA duplex variants was performed using `AlphaFold` (https://alphafoldserver.com/ accessed in December 2025) [60]. We modelled the complex in a protein molecule:DNA duplex ratio of 1:1 as our biochemical and crystallographic studies had confirmed such a binding model before. First, the modelling parameters were optimised by comparing the generated model of the NKX2.1 binding domain and the 12 bp DNA fragment containing the standard binding motif with the previously unpublished crystal structure at a resolution of 1.5Å. The parameters were further validated by generating five additional complex models and comparing them with unpublished complex crystal structures featuring single-point mutations within the binding motif of the dsDNA; the root mean square deviation (RMSD) of such comparisons ranged from 0.7 to 1.1Å. This involved calculating the RMSD of backbone atoms between the predicted and experimentally derived structures, along with a qualitative assessment of key protein:DNA interface interactions. This comparative analysis provided a basis for assessing the reliability of predictions for novel DNA variant complexes.

Selected parameters were used to generate models for complexes of the NKX2.1 binding domain with variant 12 bp dsDNA constructs, for which the crystal structures had not been obtained experimentally. For each DNA variant, the corresponding sequence was provided alongside the NKX2.1 protein sequence as input. The resulting models were evaluated based on the confidence metrics inherent to `AlphaFold`, including the *predicted local distance difference test* (pLDDT) for individual protein and DNA regions, and the *interface predicted template modeling* score (ipTM) for assessing the accuracy of intermolecular contacts.

### Preparation of double-stranded DNA for electromobility shift assays

Single-stranded DNA (ssDNA) molecules for controls and mutant libraries were synthesised by Integrated DNA Technologies (Leuven, Belgium) with equal ratios of nucleotide incorporation at sites, where variation was desired (25:25:25:25 of A:C:T:G). Three mutant libraries (**CORE**, **FLANK**, **ALL**) were designed using the *NKX2-1* binding site at the thyroglobulin promoter sequence, which was also used for MST. The specific sequences were flanked by standard Illumina adapter sequences. The libraries were named according to the regions of the thyroglobulin promoter where the variations had been introduced (for full sequences used please see **Supplementary Table S1: oligos 3–5**). The **CORE** library includes variation in the core CAAG sequence of the thyroglobulin promoter binding site (4 bp total). The **FLANK** library includes variation in the 5’ flanking nucleotides upstream and downstream of the core sequence (10 bp total) with the core sequence (CAAG) kept constant. The **ALL** library includes variation in both the flanking and the core sequences (14 bp total). The different libraries allowed us to investigate the effect of variation of the core and flanking sequences independently, while also including an even less biased design for the **ALL** library. We used positive (PC) and negative controls (NC) to optimise the EMSA conditions. The PC was the unvaried thyroglobulin promoter sequence and the NC was a length-matched exonic sequence of the *FOXA1* gene that was negative for NKX2.1 binding (for full sequences used please see **Supplementary Table S1: oligos 6–7**).

Double-stranded DNA was produced using a Klenow enzyme with a reverse primer-only, specific to the mutant libraries (**Supplementary Table S1: oligo 8**), the negative control (**oligo 10**), or the positive control (**oligo 12**). ssDNA template was incubated with 0.2 mM each of dATP, dCTP, dTTP and dGTP, 12 µM reverse primer either unlabelled or labelled with Alexa Fluor 647 dye in 1X NEBuffer 2 (New England Biolabs, Frankfurt, Germany) at 95°C for 5 minutes and slowly cooled to room temperature before adding 10 units of DNA Polymerase I, Large (Klenow) Fragment (New England Biolabs) and incubating at 20°C for 15 minutes. dsDNA was then purified using phenol-chloroform-isoamyl precipitation and the size and purity was controlled using agarose gel electrophoresis.

### Electromobility shift assays

Electromobility shift assays were performed in 8% 29:1 BIS/acrylamide, 0.5x Tris-Borate-Ethylenediaminetetraacetic acid (TBE), pH 8.2 gels. Gels were pre-run in the Mini-PROTEAN^TM^ Tetra Cell (Bio-Rad, Feldkirchen, Germany) at 100 V for at least 1 hour at 4°C. Binding reactions were performed with 2.3 pmol of GFP-NKX2.1 protein, 4.6 pmol Alexa Fluor 647-labelled mutant library dsDNA (**CORE**, **FLANK**, **ALL**), with added 0.46 fmol of unlabelled PC and NC and in a final volume of 20 µl of binding buffer [250 mM HEPES pH 7.5, 1.85 M NaCl, 10% propanediol, 0.5% Tween 20, 0.1 µg/µL poly(dI.dC), 67 ng/µl bovine serum albumin, 0.05% Nonidet P-40, 4% Ficoll400]. A no-protein control was used in every EMSA run to visualise the migration of the unbound DNA which was then extracted and sequenced to serve as a reference for the downstream enrichment analysis of bound sequences. We tested a range of DNA:protein molar ratios (from 1:10 to 1:0.1) in the binding reactions to determine the best ratios for the detection of specific NKX2.1:DNA binding (data not shown). Higher DNA:protein ratios led to equal binding of PC and NC sequences. Decreasing the DNA:protein ratios resulted in more specific binding, but the ratio should not be so low that only very strong binding sites would be detected. It is important to optimise this ratio specific to the TF under investigation. Quantitative polymerase (qPCR) reactions were then used to control binding reactions for binding specificity before performing next generation sequencing. Reactions were incubated at 30°C for 20 minutes before the addition of dsDNA, then incubated for a further 45 minutes at 30°C. Reactions were incubated for 5 minutes on ice, Orange G dye was added to a final concentration of 0.0002% w/v and then loaded onto the EMSA gel, which was run at 100 V for 1 hour at 4°C in the dark. Gels were imaged with the CHEMI® Premium Imager (VWR, Dresden, Germany) for 30 seconds using 635 nm (red) excitation and a 705 nm filter to capture the fluorescence of the Alexa Fluor 647-labelled dsDNA and using 470 nm (blue) excitation and a 525 nm filter to capture the fluorescence of the GFP-labelled NKX2.1 protein. To enhance visualisation of the protein-bound DNA bands, the maximum displayed pixel intensity was adjusted to 100% using ImageJ software, while the gamma value was kept unaltered.

### DNA purification and sequencing library preparation

DNA bands of interest were excised from the gel using clean scalpels and immediately subjected to DNA electroelution using the Model 422 Electro-Eluter (Bio-Rad) with 12–15 kDa molecular weight limiting membrane caps according to the manufacturer’s instructions. The electroelution was performed in 0.5x TBE, pH 8.2 buffer at 10 mA/gel piece for 30 min. Leads were reversed for 1 minute at 10 mA/gel piece to remove any DNA adhering to the membrane. Eluted DNA was precipitated overnight with 1 volume isopropanol, 1/10 volume 3 M sodium acetate, pH 5.2 and 200 µg glycogen, then centrifuged for 30 minutes at 4°C at 14,000 *x g*. The isopropanol was removed and the DNA pellet rinsed with 150 µl cold 100% ethanol, centrifuging again for 1 minute at 4°C at 14,000 *x g*. On removing ethanol, the pellet was allowed to air dry for 5 minutes before being resuspended in 10 µl Tris-Ethylenediaminetetraacetic acid (TE) buffer and the concentration of DNA being measured.

For Illumina sequencing library preparation, approximately 1 ng of electro-eluted DNA was used as template in a barcoding reaction with **oligos 13** and **14** (**Supplementary Table S1**) with NEBNext® High-Fidelity mastermix (New England Biolabs). Uniquely barcoded reverse primers were used to label different samples to allow for pooling prior to sequencing on the same flow cell. The thermocycler ramping rate was reduced to a maximum of 3°C/sec and the number of PCR cycles was kept low at 10-14 cycles to minimise self-annealing of non-complementary strands. We used 2.5x sample volume of the AMPure® XP Reagent (Beckman Coulter, Krefeld, Germany) for purification, performed size selection of 140 bp, and washed the library twice with 100% ethanol before eluting the DNA in 25 µl of 10 mM TRIS-hydrochloride at pH 8.0. The size and purity of libraries was assessed both by the Agilent 2100 BioAnalyzer (Agilent, Waldbronn, Germany) and by quantitative polymerase chain reaction (qPCR). The libraries were sequenced on different Illumina sequencing platforms depending on the desired read depth. The Illumina sequencing read and index primers are found in **Supplementary Table S1: oligos 15** and **16**.

### Quantitative polymerase chain reaction

qPCR was performed to assess the relative amounts of positive (**Supplementary Table S1: oligos 11 and 12**) and negative controls (**Supplementary Table S1: oligos 9 and 10**) that had been spiked into the binding reactions at a ratio of 1:10,000 to the mutant libraries. qPCR was performed in triplicates on the qTower3® thermocycler (Analytik Jena, Jena, Germany). The reactions, with a total volume of 20 µl, contained 1 ng of template DNA, 1x SYBR Green® Master Mix (Thermo Fisher Scientific) and 0.1 mM of each primer. The qPCRsoft4.1 software (Analytik Jena) was used for fluorescence quantification and cycle threshold (C_t_) calculation.

### Processing and analysis of EMSA-seq data

Ensuing `*.fastq` files from Illumina sequencing were first trimmed to 40 bp in length to remove indices and unspecific nucleotides and filtered to remove low-quality fragments with an average Phred score <30 using `cutadapt` [61]. A second trimming step removed all nucleotides except the 14 nucleotides of the regions that were carrying the randomised nucleotide positions across all three libraries. For the **CORE** libraries, an additional step of `cutadapt` processing was performed to ensure that the sequences flanking the core motif matched the wildtype sequence and the **FLANK** libraries were processed to ensure the presence of the core motif (CAAG).

The occurrence of each unique sequence was counted for the individual samples. The **FLANK** and **ALL** library samples contained high numbers of sequences matching the **CORE** library (identical flanking regions to the four randomised core nucleotides). This contamination may have occurred during dsDNA preparation (before the binding experiment) or during NGS library preparation due to barcode hopping (after the binding experiment). To prevent skewing of the binding results, we removed these fragments prior to analysis using `cutadapt`.

`DESeq2` v1.50.0 [62] was used to determine log2 fold enrichment or depletion of DNA sequences in the mutant library upon protein binding. Prior to `DESeq2` analysis, LFCsamples were filtered to remove sequences with less than five occurrences in the unbound library to avoid inaccurate and skewed LFC calculations due to low sequence counts. Experimental reproducibility was assessed using Euclidean sample-to-sample distance calculations and principal component analysis. These analyses were conducted using count data that had been normalised and subjected to a regular log transformation. `DESeq2` was run on each mutant library separately, with the unbound, 1:0.2 ratio, and 1:0.1 ratio as distinct conditions. The log2 fold change (LFC) of each DNA sequence was calculated for both 1:0.2 and 1:0.1 ratios for bound *versus* unbound fragments. Empirical Bayes shrinkage of LFC calculations was performed with the `aplegm` package to reduce noise of DNA sequences with low counts or high variability [63]. Prior to the training of deep learning models, the separate LFCs (1:0.1:unbound and 1:0.2:unbound) for each mutant library were combined by calculating the mean LFC across both conditions.

### Neural network model training

The mean LFCs were calculated using `DESeq2` and were standardised and used to train artificial neural network models, resulting in one model per mutant library. Prior to model input, sequences were processed to reconstruct the local genomic context and account for strand invariance. To represent the full promoter motif, the 14 bp variable region was padded with fixed sequences corresponding to the flanking wildtype thyroglobulin promoter sequence (5’-CACTGC…………..TTGA-3’), resulting in 24 bp sequences. We trained the model once with and once without inclusion of the respective reverse complement sequences. Comparison revealed that the model performed better if reverse complement sequences were included in the training data (data not shown). The reverse strand sequences were generated and assigned the identical LFCs as calculated for the forward strand. These additional sequences were concatenated with the original dataset, thereby doubling the amount of training data and allowing the model to learn sequence patterns independent of strand orientation. This is representative of the true biomolecular situation where TFs interact with both DNA strands of the double helix. Prior to training, the input data was standardised to zero mean and unit variance to deal with differing scales and ensure efficient convergence.

Several network architecture types were compared during training based on their performance on a hold-out validation dataset (20% of the remaining data after removing a separate 20% test set). We evaluated a diverse set of models, including multilayer perceptrons as a starting point for the model design, recurrent neural networks for capturing sequential dependencies, and equivariant networks designed to explicitly encode strand invariance. Additionally, we tested standard one-dimensional convolutional neural networks (CNNs) and multi-input models capable of combining sequence data with other numerical features. From these candidates we chose a CNN architecture that combines dilated convolutions, as seen in models like BPNet [64], and adaptive max pooling [65] because it performed the best across all mutant libraries.

The model was trained to predict the LFC using the one-hot encoded sequence as input, meaning the DNA sequence is converted into a binary vector [66]. Although the models were trained with 24 bp long sequences, they can be used to predict the LFC of shorter or longer DNA sequences. In this project we tested lengths of 12, 14, 17, 24 and 500 bp. We optimised the hyperparameter settings for each model independently, using the settings that produced the least mean-squared error. The software packages used are listed on **Supplementary Table S2**. The final hyperparameter settings can be found in **Supplementary Table S3**.

### ChIP-seq validation

The performance of each model was validated using as ground truth for the accuracy of TFBS predictions various publicly available ChIP-seq data sets (**Supplementary Table 4**). To test the models’ ability to distinguish between competing biological signals, we employed them to classify ChIP-seq peak sequences from experiments for NKX2.1 (positive dataset) and for other TFs (negative datasets). As negative control TFs, we used GATA1, MYOD1, NKX2.5 and RXRA. All datasets were acquired from ChIP-Atlas and were filtered for high-confidence peaks (MACS2 *Q*-value <10^-50^). We trimmed the ChIP-seq peaks to a length of 500 bp around the peak centre. Any overlapping genomic regions between the positive and negative datasets were removed using `bedtools` [67] version 2.30.0. We used our models to predict the LFC of the 500 bp sequences from both positive and negative control TF ChIP-seq experiments. The LFC prediction was then used to classify the peak as belonging to the NKX2.1 dataset or the negative control TF dataset. We used *Find Individual Motif Occurrences* (`FIMO`) v5.5.7 [68] with the mouse Nkx2.1 PWM from JASPAR (2024 release, MA1994.1) [69] for the same task as a performance baseline. The PWM was slid across the 500 bp sequence and the maximum scoring sequence was used to represent each peak sequence. Since `FIMO` relies on traditional PWMs, it serves as a benchmark to quantify the predictive improvement offered by the neural network’s ability to process non-linear sequence dependencies. In addition to using the whole 500 bp sequence for model prediction, we tested a sliding window approach, similar to the use of the PWM model. Since training sequences were 24 bp long, we used a sliding window of this length across the ChIP-seq peak sequences. We then took the 24-mer with the highest LFC to represent the peak.

### Positional bias analysis

We then asked where in the ChIP-seq peak sequences the models with a 24 bp sliding window predicted the strongest binding sequence. The sequence with the highest LFC was extracted for each peak and the first nucleotide of this 24 bp sequence was used to reference its position. This was repeated using the PWM. The resulting distributions were visualised as normalised histograms to represent position occurrence in percent (**Figure 11**).

## Results and Discussion

### EMSA-seq is able to quantify the effects of DNA sequence variation on transcription factor binding in a high-throughput manner

Electromobility shift assay sequencing (EMSA-seq) is a method we adapted from Wong *et al.* [16], Gu *et al.* [17] and Levo *et al.* [18], to quantify the enrichment of sequences in a mutant library upon TF binding. Sequencing the mutant library before and after binding allowed us to quantify the enrichment of each sequence relative to the reference unbound library. Unlike HT-SELEX, EMSA-seq uses only one round of TF-binding purification and is therefore able to detect low levels of enrichment, or low affinity binding sites. It has been shown that low affinity binding sites are important for normal development [70,71], and even small changes in binding affinity can disrupt developmental processes [5,72]. Using EMSA-seq, we investigated the relative binding of NKX2.1 to millions of DNA sequences in parallel.

We designed three mutant libraries based on the NKX2.1 binding site in the rat thyroglobulin promoter [59]. The libraries differ in the region where variation was introduced. The **CORE** library contains variation in the four central nucleotides, the **FLANK** library contains variation in the sequence flanking the core nucleotides (five nucleotides on each side), and the **ALL** library contains variation in the entire 14 bp sequence that includes the core and flanking nucleotides. An example for the EMSA of the **CORE** library can be seen in **Figure 1A**. qPCR was performed on each sample using primers specific for the positive (PC) and negative (NC) controls. The average cycle threshold values (C_t_) are displayed in **Figure 1A**. In both lanes, the PC was enriched by a factor of ∼40 fold indicating that these conditions allowed for specific DNA binding by NKX2.1. We tested a range of molar ratios in the binding reactions but found that an increase in the protein concentration resulted in higher variability between the replicates and unspecific binding (data not shown). This highlights the importance of testing a range of conditions for the binding reaction in order to secure specific binding with EMSA-seq. Probably, such optimisation has to be performed independently for each unique TF due to variability in binding affinities along with other variables such as temperature, incubation time and buffer composition. `DESeq2` was used to calculate the log2 fold change (LFC) of DNA sequences in the bound *versus* unbound conditions. `DESeq2` is traditionally used for gene expression analysis but can be applied to other genomic count data that follows a negative binomial distribution, as is the case for EMSA-seq data [62]. The Minus-Average (MA) plots in **Figure 1** show the LFC calculations for DNA sequences in the **CORE** library across the two DNA:protein molar ratios. The wildtype CAAG core sequence was the most enriched in both conditions. The **FLANK** and **ALL** library EMSA results are found in **Supplementary Figure S1** and the full `DESeq2` results in the GEO repository (GSE319141).

**Figure 1:**
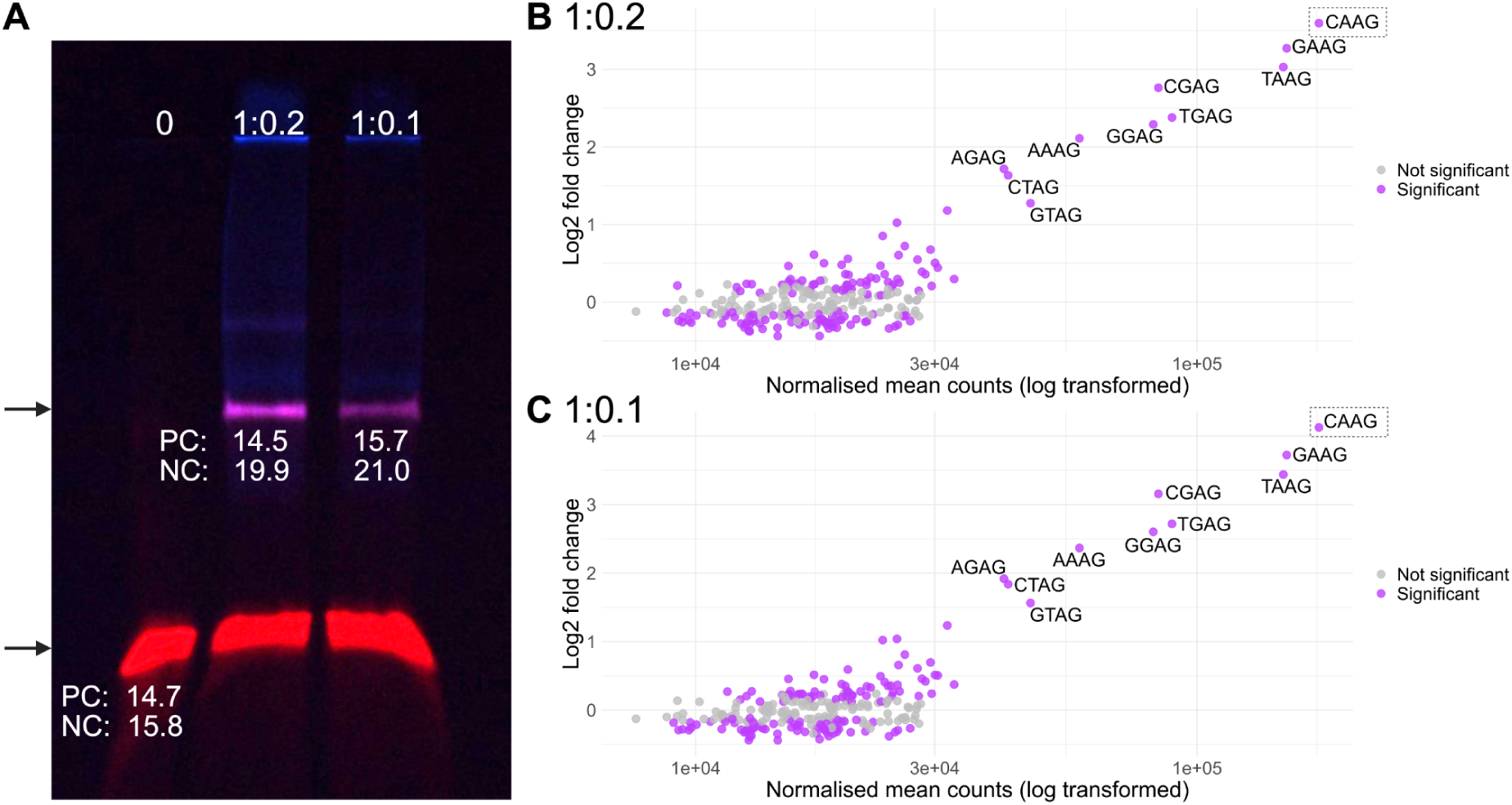
**A:** Electromobility shift assay with NKX2.1-GFP (blue) and the **CORE** library DNA (red). The DNA:protein molar ratios of each lane are indicated at the top (0, no protein control). The top arrow indicates NKX2.1-bound DNA and the lower arrow indicates unbound DNA. The extracted DNA was used in a qPCR reaction with either positive control- (PC) or negative control- (NC) specific primers. The reactions were performed in triplicate and the average cycle threshold (C_t_) value for each PC and NC primer pair is indicated below the corresponding bands. **B and C:** MA plots of the `DESeq2` results comparing the log-transformed sequence counts *versus* the mean `DESeq2`-calculated log2 fold change for the 1:0.2 ratio (**B**) and the 1:0.1 ratio (**C**). The data points are coloured according to significance (purple, *padj* < 0.1, grey, n.s.). The ten sequences with the highest calculated log2 fold change are labelled with their respective core sequence. The wildtype sequence is most enriched in both molar ratios (dashed box).

EMSA-seq is a relatively simple, inexpensive method that uses commonplace laboratory equipment. We used fluorescently-tagged DNA and protein in our binding experiments, which in theory could sterically affect TF binding preferences. It is possible to use unlabelled protein, however, we wanted to ensure comparability with MST, which requires a fluorescent label for molecule tracking. We used a single-end DNA label that was separated by a constant 24 bp primer-binding sequence, rendering it unlikely to impede NKX2.1 binding. Additionally, the specificity of binding reactions was assessed with unlabelled positive and negative DNA controls. EMSA-seq could be performed with entirely unlabelled molecules, using labelled DNA and/or protein in neighbouring EMSA wells to serve as a reference for DNA extraction.

We only used the DNA binding domain (DBD) of NKX2.1 for our experiments and it is possible that the full length protein might have resulted in different binding preferences. However, Jolma *et al.* [20] discovered that the DBD was primarily responsible for TF-DNA binding specificities for all but one (ELK1) of the hundreds of the human TFs studied. It has been reported that the Nkx2.1 N-terminal contacts the DNA minor groove, but this is thought to have a role in transactivation rather than DNA-binding specificity [73]. Phosphorylation of the NKX2.1 protein has been implicated in differential gene regulation of specific promoters [74–79]. Future studies could investigate the effect of post-translational modifications using EMSA-seq with modified and unmodified versions of the TF.

### EMSA-seq-derived convolutional neural network models learn TF binding specificities

The `DESeq2` results from the EMSA-seq experiments were used to train deep learning models to learn the NKX2.1:DNA binding specificities and predict the LFCs of DNA sequences not captured by the EMSA-seq experiments. It is important to be aware that the LFC is a unitless measure of enrichment associated with NKX2.1 binding, rather than a direct assessment of DNA:protein binding affinity (e.g., its free energy). Importantly, the LFC is relative and thus comparable only between DNA sequences of identical length. Several model architecture types were tested for their ability to predict the LFCs of hold-out validation sequences (20% of the data without the test dataset). The model that had the lowest validation loss across mutant library types was a combination of BPNet [64] and variable convolutional neural network (VCNN) architectures.

We introduce VCNNBPnet, a deep CNN designed for the analysis of histone-free linear DNA sequences (**Figure 2**). The architecture was adapted from the BPNet framework [64] but optimised for sequence-level scalar predictions rather than base-pair resolution profiling, resulting in a score per sequence instead of per nucleotide. The model takes one-hot encoded DNA sequences as input and processes them through a stack of dilated convolutions to capture multi-scale motif interactions, followed by a global pooling mechanism and a dense block for final prediction. The input to the model is a tensor 𝑋ɛ𝑅^𝐵×4×𝐿^ , where *B* represents the batch size, and *L* represents the sequence length. The four input channels correspond to the one-hot encoded nucleotides (A, C, G, T). The core feature extractor consists of a series of 1D convolutional layers. We used dilated convolutions to efficiently expand the receptive field without downsampling or increasing the parameter count. The network sequentially applies a set of dilation rates, e.g. D=[1, 1, 2, 4, 8]. This hierarchy allows early layers to capture local sequence motifs (such as individual TFBSs), while deeper layers with higher dilation rates integrate information across wider genomic contexts to detect motif syntax and periodicity. For a kernel size (*K*) and dilation rate (*d*), we apply padding (*P*) to maintain the spatial dimension of the input sequence throughout the convolutional stack. The padding is calculated as:

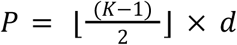

**Figure 2:**
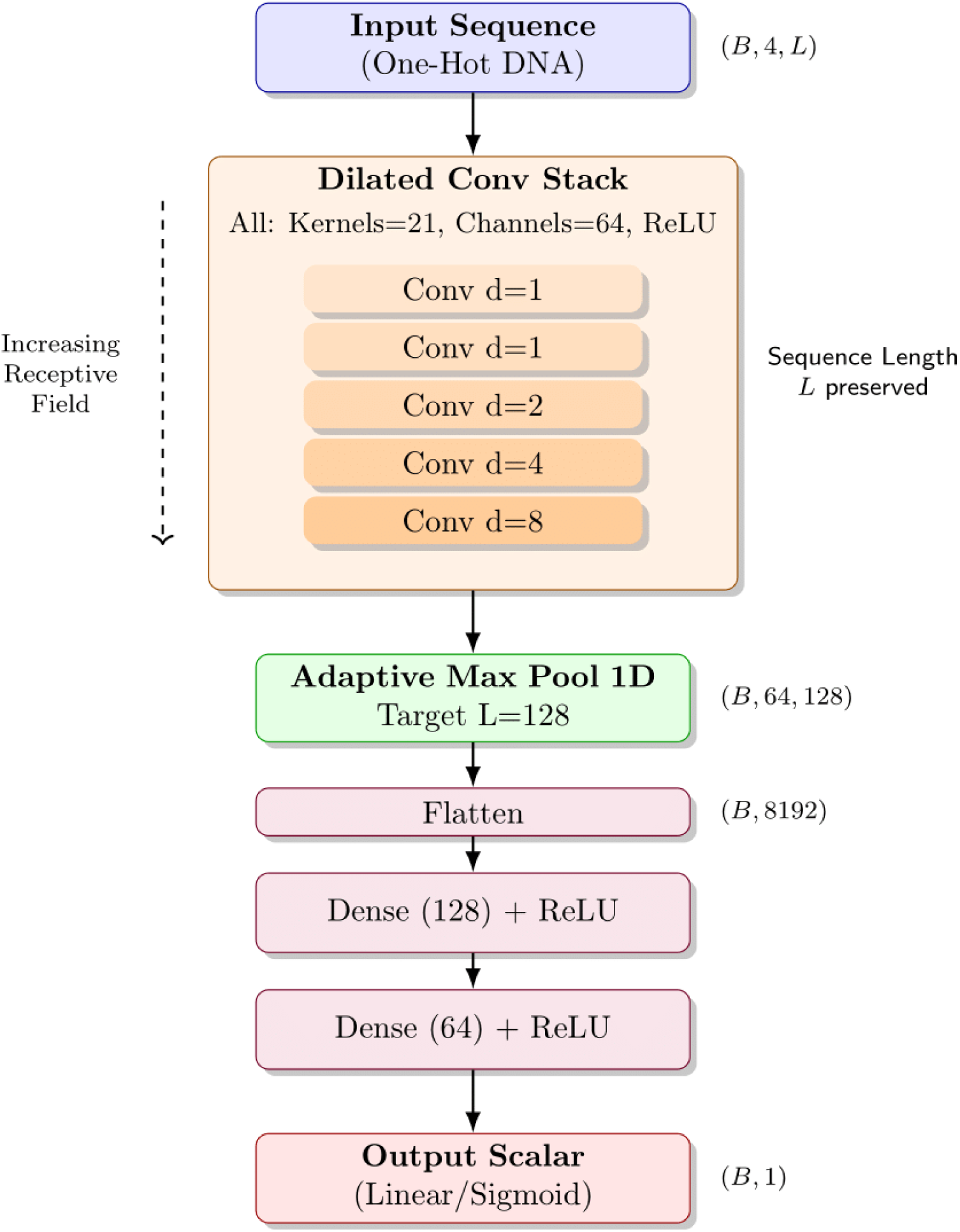
A schematic overview of the proposed VCNNBPNet framework for DNA sequence analysis. The model accepts variable-length, one-hot encoded DNA sequences (tensor shape B,4,L) and processes them through a backbone of five stacked convolutional layers. This stack uses 64 filters with a kernel size of 21 and employs exponentially increasing dilation rates (d=1,1,2,4,8) to expand the receptive field and capture distal regulatory motifs without reducing input resolution. To standardise feature dimensions across varying input lengths, feature maps are downsampled *via* an adaptive max pooling layer to a fixed target length of 128. The resulting representation is flattened and passed through a dense block, comprising two fully connected layers (128 and 64 units), with ReLU activation to generate the final scalar prediction. This graph shows an example architecture; hyperparameters can differ for the different models.

Each convolutional layer is followed by a rectified linear unit (ReLU) activation function. Following the convolutional encoder, the feature maps are passed to an adaptive max pooling layer. Unlike standard max pooling, which uses fixed window sizes, this layer forces the variable-length spatial dimension into a fixed output size 𝑆𝑝𝑜𝑜𝑙 (default 128).

This decouples the architecture from specific input lengths, allowing the model to handle variable-length DNA fragments. The pooled features are flattened and processed by a multi-layer perceptron. This dense block consists of fully connected layers with decreasing hidden sizes (e.g., 128-64), utilising ReLU activations.

The final output layer projects the latent representation to a scalar value. Depending on the task (classification *versus* regression), a sigmoid activation or linear pass is applied to predict binary binding probability or continuous signal strength, respectively. Additionally, VCNNBPNet utilises a stacked convolutional layer architecture rather than the residual (ResNet) blocks found in the original BPNet, favouring architectural simplicity for datasets where ultra-deep networks may risk overfitting. The inclusion of an adaptive pooling layer replaces the profile prediction head, acting as a bottleneck to summarise motif presence across the entire input window.

We trained one model for each mutant library, using the average EMSA-seq LFC across the two DNA:protein molar ratios (1:0.2 and 1:0.1) as input. The correlation between the molar ratios was high for **CORE** and **FLANK** libraries (Spearman coefficients of 0.98 and 0.85, respectively), while the **ALL** library correlation was slightly lower (Spearman coefficient = 0.69). The sequences with the highest variability were those with zero counts in half of the bound samples. Subsequently, using the average LFC across the molar ratios for the **ALL** library is expected to stabilise variation caused by low sequence counts. Although the EMSA-seq experiment used DNA sequences of 70 bp, the training was performed with DNA sequences of 24 bp (the experimental sequence with Illumina sequencing adapters removed to reduce computational cost by reducing the DNA length). We used the full 24 bp promoter sequence, despite only a maximum variable range of 14 bp to capture any interdependencies between nucleotides in the varied and constant regions. The models were tested for their performance on the holdout data and resulted in the mean-squared-errors of 0.0459 for the **CORE** model, 0.0514 for the **FLANK** model, and 0.2849 for the **ALL** model. This is in line with the validation loss seen during the training of each model (**Supplementary Figure S2**).

The variable results between VCNNBPNet models demonstrate that while EMSA-seq is able to assess TF binding to millions of sequences in parallel, increasing the mutant library sequence diversity requires increasingly high read depths. The **ALL** mutant library has >200 million possible unique DNA sequences. It is important to consider the trade-off between the length of the DNA to be randomised and the cost of sequencing. We found that a sequence diversity of 10 bp (**FLANK** library) was sufficient to predict relative NKX2.1:DNA binding strengths through the learning of complex patterns, as discussed below.

We used `tangermeme` [58] to perform Shapley value estimation (`DeepSHAP`) [80] and *in silico* saturation mutagenesis to assess the contribution of each nucleotide to the model’s LFC prediction. The wildtype thyroglobulin promoter’s NKX2.1 binding site sequence patterns are depicted in **Figure 3**. Shapley values measure the size of each nucleotide’s attribution in the LFC prediction and take into account the possibility of nucleotide interdependencies. In contrast, *in silico* saturation mutagenesis, LFC prediction changes are a measure of how each nucleotide affects the LFC prediction, but with only one nucleotide considered at a time, thereby neglecting the role of interdependencies. Encouragingly, the patterns across all models and both analyses are similar.

**Figure 3:**
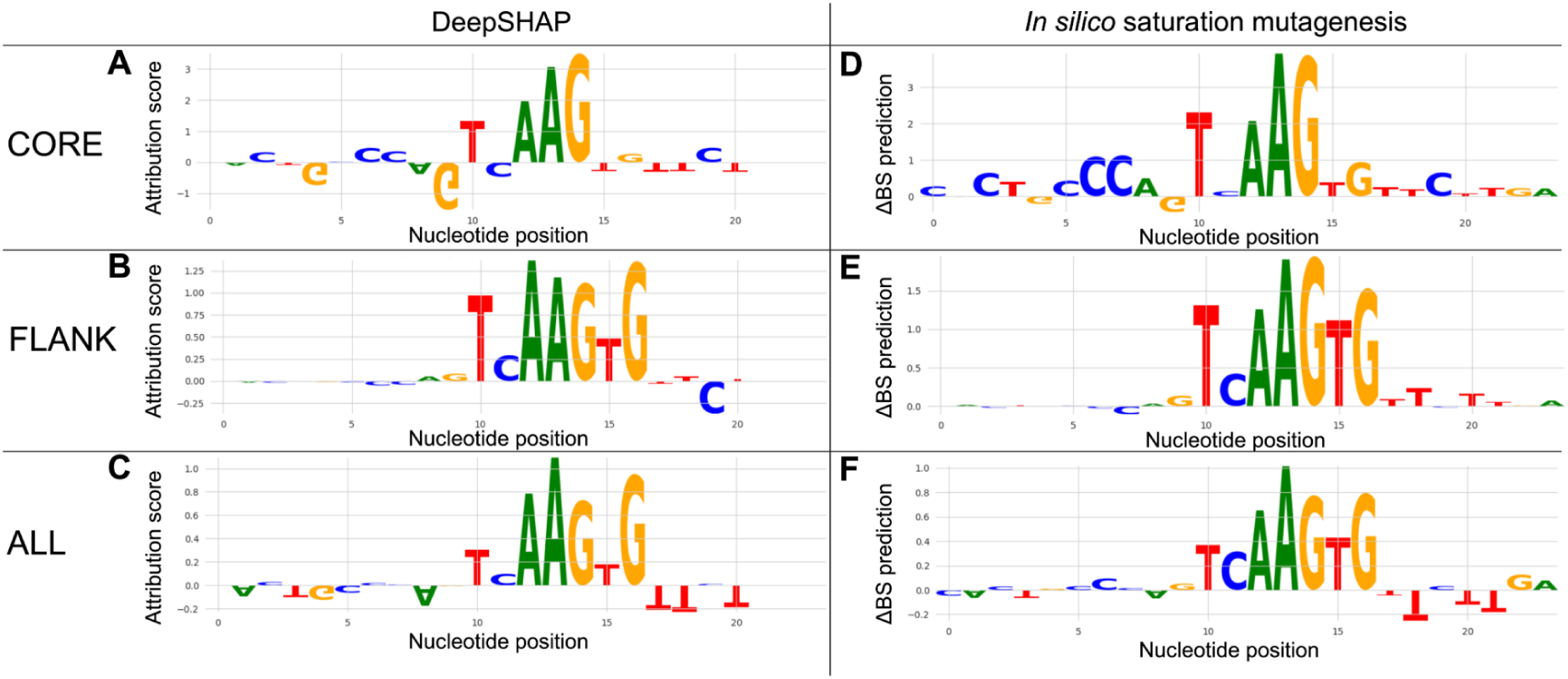
DeepSHAP attributions and *in silico* saturation mutagenesis profiles by each model for the wildtype thyroglobulin promoter NKX2.1 binding site sequence. **A: CORE**, **B: FLANK**, and **C: ALL** models Shapley values. **D: CORE**, **E: FLANK**, and **F: ALL** models *in silico* saturation mutagenesis prediction changes. The Shapley value estimations represent the attribution each nucleotide plays in the model’s LFC (BS) prediction where the height of the nucleotide represents the attribution score. The *in silico* saturation mutagenesis profile nucleotide size represents the change in the model’s LFC (ΔBS) prediction when the nucleotide is mutated.

The **CORE**, **FLANK**, and **ALL** models assigned remarkably similar importance scores to nucleotides, even in regions that were held constant during the experiment. For instance, the **FLANK** model correctly identified critical patterns in the core region, mirroring the results of the comprehensive **ALL** model. This attribution of importance to unmutated regions reflects the underlying sequence context learned by the model, which is subsequently revealed by the DeepSHAP analysis. During training, the model internalised that high LFCs depend on the constant regions as much as the variable ones. DeepSHAP elucidates this by calculating contribution scores against a randomised background, confirming that the model treats these constant regions as essential signals rather than neutral noise. Ultimately, the model has inferred a combinatorial logic, recognising that variable motifs only result in high binding affinity when supported by the specific context of the constant scaffold. Overall, the neural network learns binding motifs in a position-independent manner; it recognises that a specific sequence pattern is important regardless of whether it was mutated in a specific library.

The use of neural networks over simpler models such as PWMs, allows the detection of more complex patterns shaped by interdependencies between nucleotide positions, unlike TFFMs, which just consider neighbouring dinucleotides. The term “epistasis” refers to the nonindependence of biological units [81]. We used this concept to identify positions and incidences where the effect of a double point mutation in the NKX2.1 binding site deviated from the expectation of an additive effect of each individual mutation effect. The results from this analysis for the **FLANK** model are depicted in **Figure 4**. Interdependencies encode information on the DNA shape (e.g., bending) [82–85], which in turn affects TF binding [7,83,84,86–88]. We found that third-order dependencies exist, however the illustration of these relationships would be too complex for the 2D space (data not shown).

**Figure 4:**
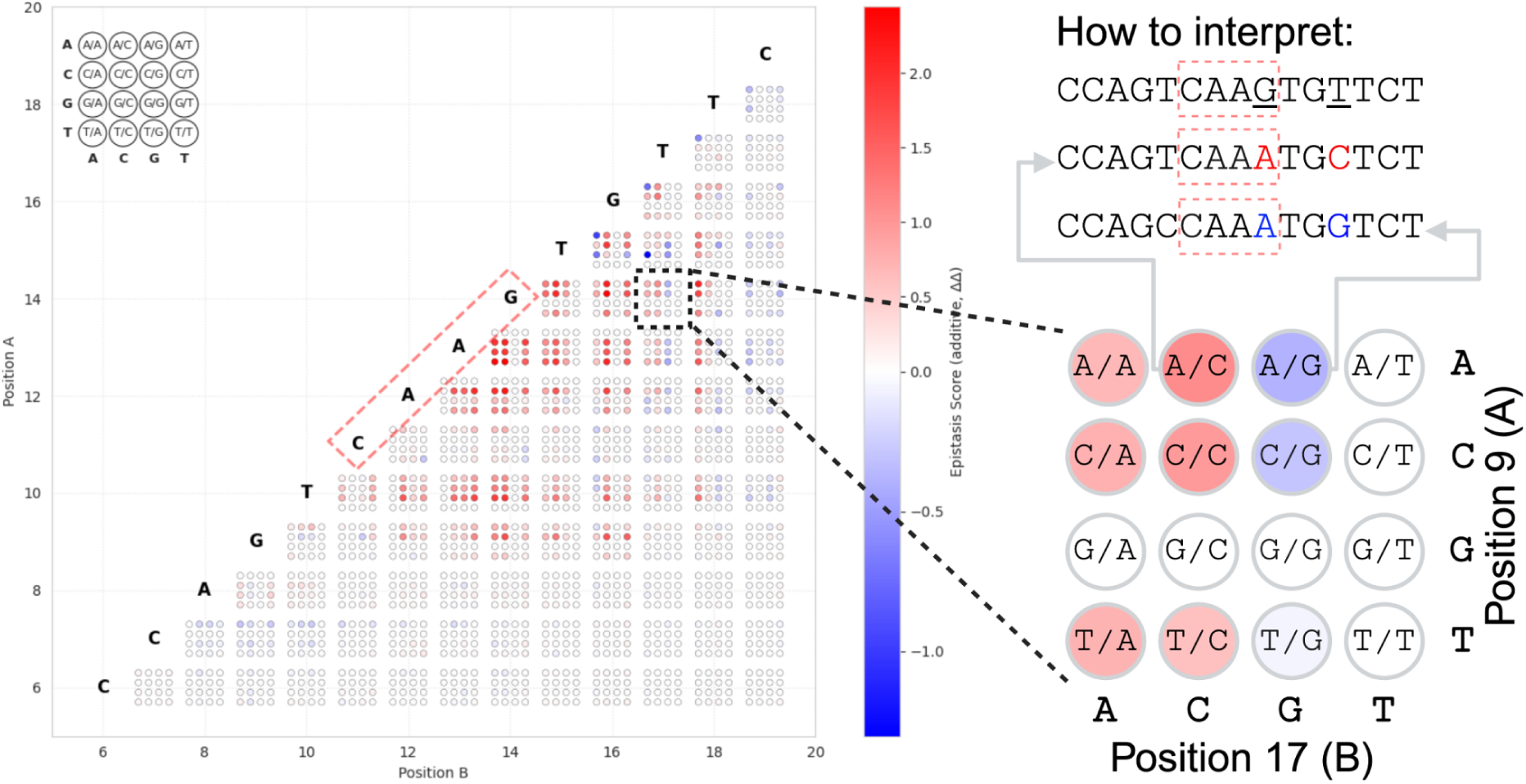
Nucleotide interdependency map of the NKX2.1 binding site for the **FLANK** model. The wildtype thyroglobulin promoter binding site sequence is shown diagonally with a red-dashed box enclosing the core nucleotides. For each double SNV, position A is on the y-axis and position B on the x-axis. The epistasis score is shown with a red-blue colour scale where a super-additive effect is red and a super-antagonistic effect is blue. The right side of the figure provides an example of how to read the map. Each square of 16 dots represents the variable combination of two nucleotide positions. Each possible nucleotide combination is depicted. The example shows that the combination G>A [at position 14] **plus** T>C [at position 17] results in increased, while the combination G>A [at position 14] **plus** T>G [at position 17] results in decreased binding compared to the expected additive effects when considering each SNV individually.

### EMSA-seq captures binding affinities at higher resolution compared to microscale thermophoresis through competitive binding dynamics

MST was performed to test the binding of NKX2.1 to n = 29 altered thyroglobulin promoter binding site DNA sequences. One or two nucleotide variants were introduced into the core sequence or the two flanking nucleotides on either side of the core sequence. The same DNA sequences were offered to each deep learning model for LFC prediction and to `FIMO` [68] using the Nkx2.1 PWM (JASPAR: MA1994.1) [69]. The Spearman correlation coefficient was calculated to compare each of the model predictions with the DNA sequence dissociation constant (*K_d_*) calculated by MST. The *K_d_* was negative-log-transformed to reflect its relationship with the change in standard Gibbs free energy on binding [89]. This analysis revealed that the *K_d_* and model predictions did not correlate (MST *versus* **CORE** r = 0.04; MST *versus* **FLANK** r = 0.00, MST *versus* **ALL** r = 0.14, and MST *versus* PWM r = -0.13). However, the predictions across deep learning model types correlated well (**CORE** *versus* **FLANK** r = 0.91, **CORE** *versus* **ALL** r = 0.84, and **FLANK** *versus* **ALL** r = 0.88), which was higher than the correlations between PWM and deep learning models (PWM *versus* **CORE** r = 0.64, PWM *versus* **FLANK** r = 0.72, and PWM *versus* **ALL** r = 0.59).

The average *K_d_* between NKX2.1 and each DNA sequence, and its predicted *relative fold change* by each model can be seen in **Figure 5**. The free binding energy (*K_d_*) and the FCs are displayed relative to the wildtype sequence so that the wildtype FC is set to 1.0 (dashed line). The majority of the DNA sequences have a *K_d_* that falls within the margin of error of the wildtype sequence (grey bar). Consistent with the correlation coefficients, no clear pattern is visible between MST data and model predictions, whilst there is a clear trend across model predictions. A possible explanation for the lack of correlation between model predictions and MST measurements could be that EMSA-seq is able to detect relative binding affinities of DNA sequences at a higher resolution than MST, perhaps due to the competitive nature of EMSA-seq. While EMSA-seq indirectly assesses TF binding to a pool of many, competing DNA molecules, MST only measures the direct binding affinity of two molecules in isolation: a single DNA sequence and its TF. Therefore, it is plausible that EMSA-seq may detect small binding affinity differences between DNA sequences that MST does not detect due to a lack of competition.

**Figure 5:**
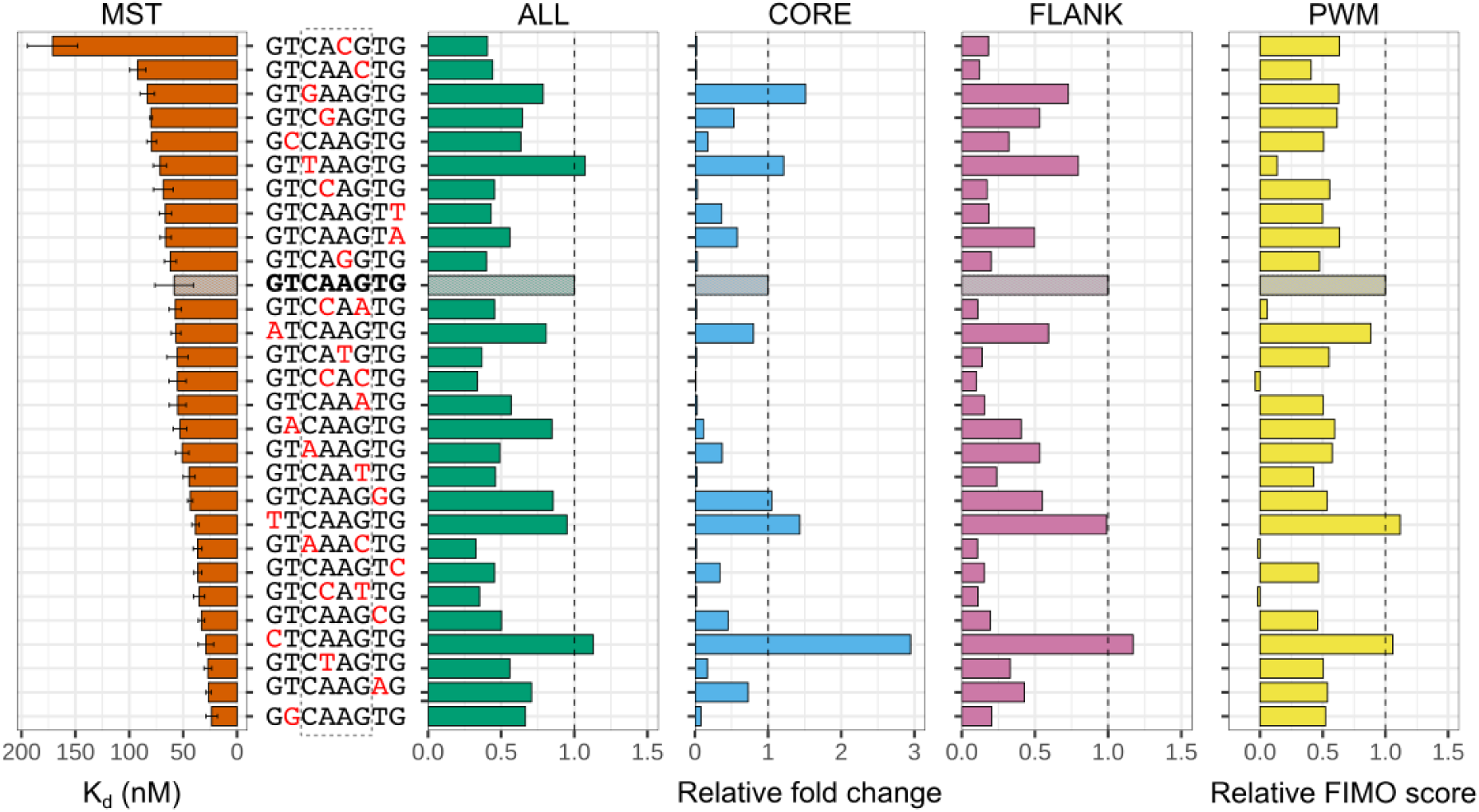
Microscale thermophoresis (MST) dissociation constants (*K_d_*) (orange) for NKX2.1 binding of n = 29 varied DNA sequences and their predicted *relative fold changes* (FC) by the **ALL** (green), **CORE** (blue), **FLANK** (pink), and PWM (yellow) models. Predicted FCs from neural network and PWM models were normalised to the wildtype score (grey bar), set to 1.0 (dashed line), called the “*relative fold changes*”. For the PWM, the highest `FIMO` score per sequence was used. The binding in MST was measured three times with the average displayed in a bar and standard deviation represented by whiskers. The sequences were 24 bp long but only the 8 bp variable region is labelled for brevity. The red nucleotides represent those that deviate from the wildtype sequence. The dashed grey box encloses the four core positions.

Fabbro *et al.* [90] performed selective chromatography (SC) with the NKX2.1 protein and n = 15 varied DNA sequences of the same wildtype thyroglobulin promoter sequence used in this study. We used our models to predict LFCs for the sequences in this experiment and found that the Spearman correlation coefficient between the experimental binding affinities and the predictions was much higher than for MST measurements (SC *versus* **CORE** model r = 0.76, SC *versus* **FLANK** model r = 0.76, SC *versus* **ALL** r = 0.77). The SC experiment was performed with all 15 DNA sequences in parallel, resulting in a competitive environment similar to EMSA-seq. This further supports the hypothesis that EMSA-seq can detect smaller binding affinity changes caused by single DNA sequence variants due to its competitive nature. Furthermore, such a competitive environment is closer to reality within the cell, where all accessible genomic regions are potentially competing for TF binding.

Discrepancies between MST and EMSA-seq binding measurements likely stem also from differences in reaction conditions, particularly temperature and the duration of incubation. While MST involves a short, low-temperature incubation followed by a thermal spike, EMSA-seq uses higher temperatures over longer periods. These thermal variations influence the biochemical properties of both molecules; e.g., increased heat enhances DNA flexibility, which may fundamentally alter TF docking [91]. Studies on p53 demonstrate that DNA binding affinity can fluctuate across temperature gradients as hydrogen bonds rearrange [92]. Furthermore, temperature modulates electrostatic and hydrophobic forces, such as the energetically favorable release of counterions at higher temperatures. [93]. Because these thermal effects are complex and highly TF-specific, they likely contribute to the observed affinity discrepancies between MST and EMSA-seq results.

Reaction timing also distinguishes MST and EMSA-seq. Recent *in vivo* data suggest that TF residence times are longer than previously assumed, typically ranging from 30 to 100 seconds [94]. While the brief MST incubation might theoretically capture TFs in a transient “sliding” search mode rather than a stable bound state, the short DNA fragments used in these specific assays likely reached equilibrium within the two-minute window.

Finally, the physical nature of EMSA-seq — specifically running the DNA:protein complex through a gel matrix — introduces unique variables. Although gel electrophoresis is sometimes thought to disrupt weak interactions, the higher resolution and broader affinity range observed here suggest otherwise. Nonetheless, the gel matrix itself imposes structural constraints on DNA:protein interactions that remain difficult to fully predict or quantify.

Next, we compared the MST binding measurements and deep learning model predictions to `FIMO` LFCs, which were calculated based on how well the input sequence matches the Nkx2.1 PWM [68]. The PWM scores mostly follow the same trend as our deep learning models, except for a couple of instances. Interestingly, for the sequence 5’-GT**CACG**TG-3’, which has very low binding in MST (*K_d_* = 171.2, SE = 23.2), the PWM has a comparatively high *relative FIMO score* of 0.63 (**Figure 5**). However, in this incidence, our models are in line with the MST measurements and predict low *relative fold changes* (**ALL** model = 0.41, **FLANK** model = 0.18, and **CORE** model = 0.02, **Figure 5**). The discrepancy between PWM and deep learning methods could be explained by the ability of deep learning to learn higher-order relationships between nucleotides, regardless of their position [8]. Conversely, PWM-based methods only consider neighbouring, dinucleotide dependencies [7]. We have used the mouse Nkx2.1 PWM (MA1994.1) from the JASPAR database [69], but the NKX2.1 DBD is 100% conserved between human and mouse, so this was unlikely to affect the performance of the PWM model.

### X-ray crystallography reveals interactions between NKX2.1 and the DNA sequence that align with patterns learnt by neural network models

Following the establishment of a robust protocol for producing highly pure NKX2.1 DBD, we initiated crystallisation experiments for high-resolution structural analysis to investigate the molecular basis of differential binding of various DNA sequences. Optimal diffraction was achieved using a 12 bp dsDNA fragment, chosen for its efficiency in yielding well-diffracting crystals, likely due to the size constraints of the NKX2.1 DBD. We obtained high resolution crystal structures including NKX2.1 DBD and the wildtype thyroglobulin promoter binding site sequence, and eight variant DNA sequences. Detailed inspection of these structures allowed us to precisely describe the intricate network of direct and water-mediated interactions between NKX2.1 and its dsDNA target.

The wildtype crystal structure (PDB ID: 9U18) demonstrates that the homeodomain adopts a canonical fold comprising an extended N-terminal loop (N-loop) followed by three α-helices (H1–H3). The N-loop anchors within the DNA minor groove, while the recognition helix (H3) resides in the major groove (**Figure 6A**). Within the N-loop, R162 and R165 facilitate direct base-specific interactions with the core CAAG binding motif and the immediately preceding position. Conversely, while H3 side chains primarily contact the phosphodiester backbone, Q210 and N211 engage in water-mediated interactions with the bases of the reverse complementary strand. We used the **FLANK** model only to show sequence attributions here, since the LFC predictions across models were consistently-well correlated. Interestingly, the nucleotides that interact with the protein were considered important by the deep learning models (**Figure 6B**), confirming their biological relevance, but the VCNNBPNet model did not discern whether a nucleotide was interacting directly or indirectly with NKX2.1. However, the more distal interacting nucleotides (at the 3’ end) were not found to be important for the model’s LFC prediction.

**Figure 6:**
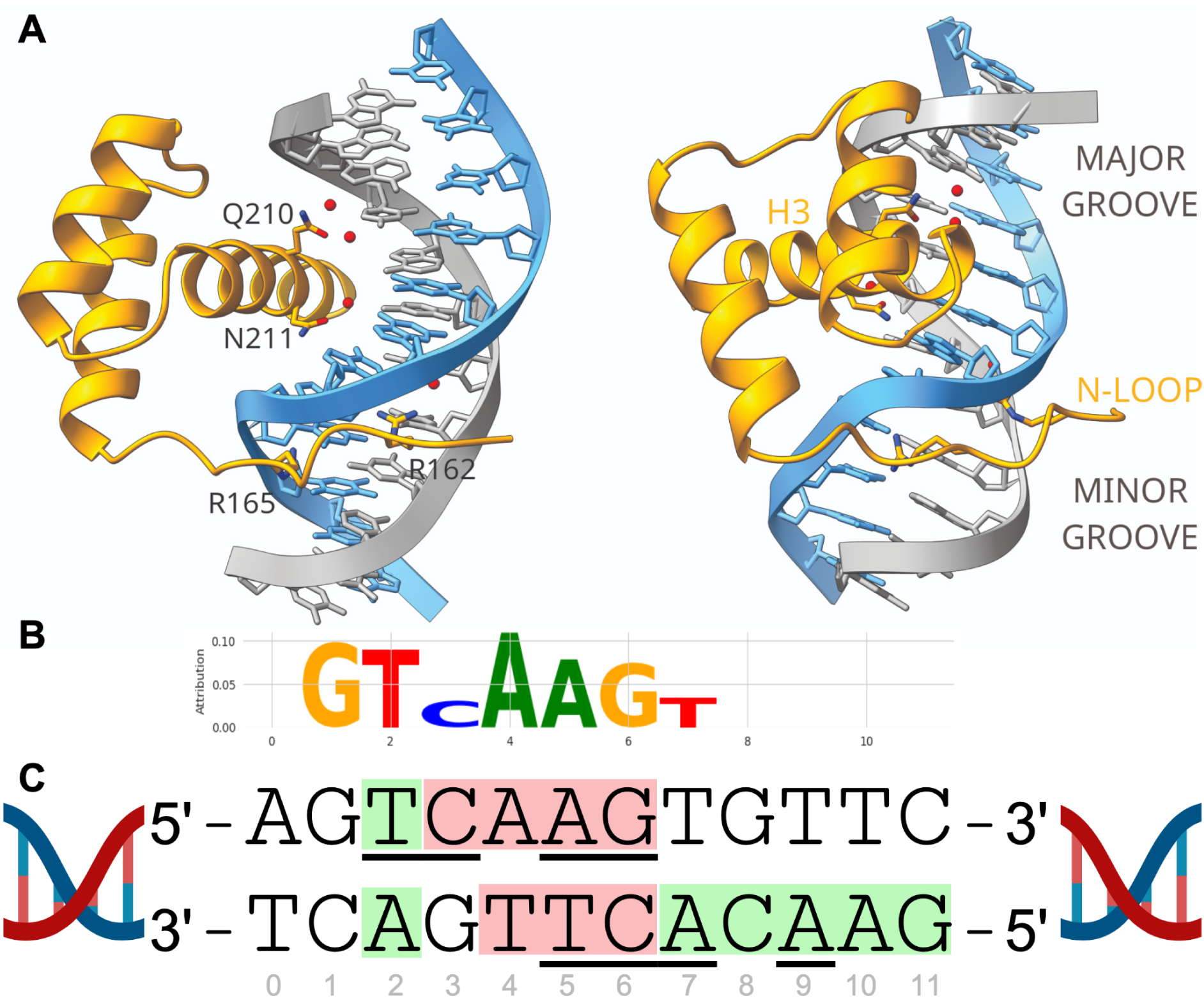
X-ray crystallography structural analysis validates patterns of attributions learnt by deep learning models. **A:** X-ray crystallography structure of NKX2.1 DBD bound to the 12 bp sequence of the thyroglobulin promoter. **B:** DeepSHAP nucleotide attributions of the thyroglobulin promoter binding site by the **FLANK** model, where size indicates importance for model prediction. **C:** Simplified diagram explaining which nucleotides interact with the NKX2.1 DBD. ***Red***: nucleotides in the core sequence that interact with the protein, ***Green***: flanking nucleotides that interact with the protein, ***Underlined***: nucleotides that interact directly with the protein (other interactions occur via the DNA backbone with either sugar or phosphate moieties). Created in https://BioRender.com.

Comparative analysis across all nine NKX2.1:DNA variant structures reveals a critical mechanistic insight: only the **CAAG>CACG** mutation (PDB ID: 9U19) induces a significant rearrangement within the protein:DNA interface (**Figure 7**). This transition is driven by a subtle shift (≤ 0.5Å) in the R165 side chain, which displaces the N-terminal loop outward from the minor groove. This movement triggers a reconfiguration of the internal hydrogen-bonding network within the protein scaffold, ultimately stabilising an alternative conformer. This structural plasticity provides a direct rationale for the altered binding characteristics observed in MST and EMSA-seq (**Figure 5**).

**Figure 7:**
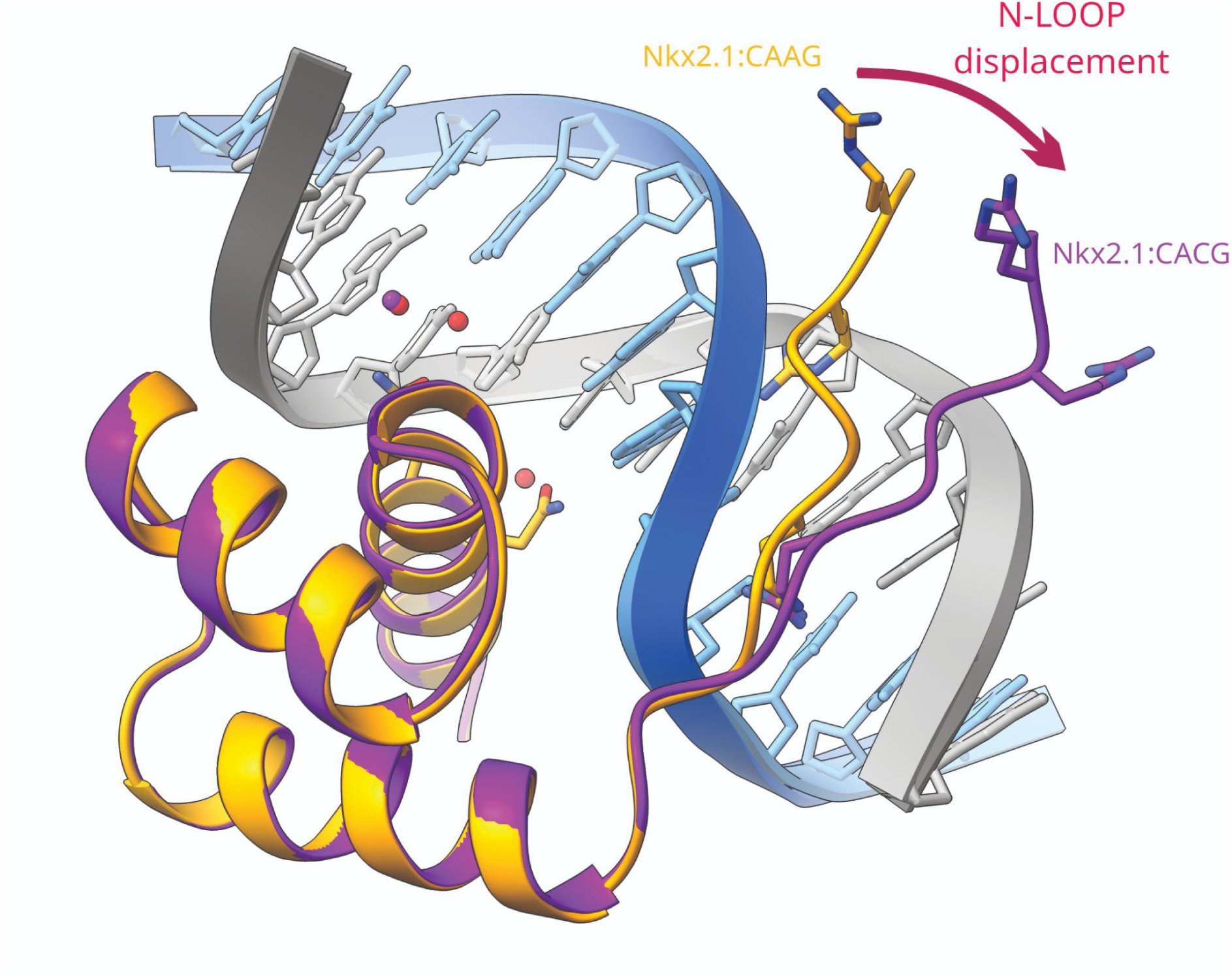
X-ray crystallography structural analysis of the CACG mutation validates patterns of attributions learnt by deep learning models. **A:** Comparison of crystal structures of the wildtype core sequence: CAAG and the SNV: CACG interacting with the NKX2.1 DBD.

Given the extensive experimental and analytical work required for X-ray crystallography, we used the above structures (PDB IDs: 9U18 and 9U19) to validate `AlphaFold` predictions for unsolved NKX2.1:DNA sequence conformations. Structural models generated by `AlphaFold` provide insight into the altered binding mechanisms of non-crystallising variants. Specifically, we investigated the role of the thymine nucleotide directly upstream of the core sequence, since its substitution with a guanine had resulted in stronger binding affinities in the MST experiment (**Figure 5**). However, the structural prediction in the presence and absence of this and other mutations (**Figure 8**) showed that the thymine to guanine substitution disrupted the local contact of thymine to Q210. This loss of interaction triggers a conformational shift in the Q210 and N165 side chains, accompanied by a slight outward displacement of the N-loop, which would result in a weaker binding between the mutated DNA sequence and the NKX2.1 DBD. This change was mirrored by a clear reduction in the LFC for this sequence in the **FLANK** model predictions which stands in stark contrast to the MST measurement (**Figure 5**).

**Figure 8:**
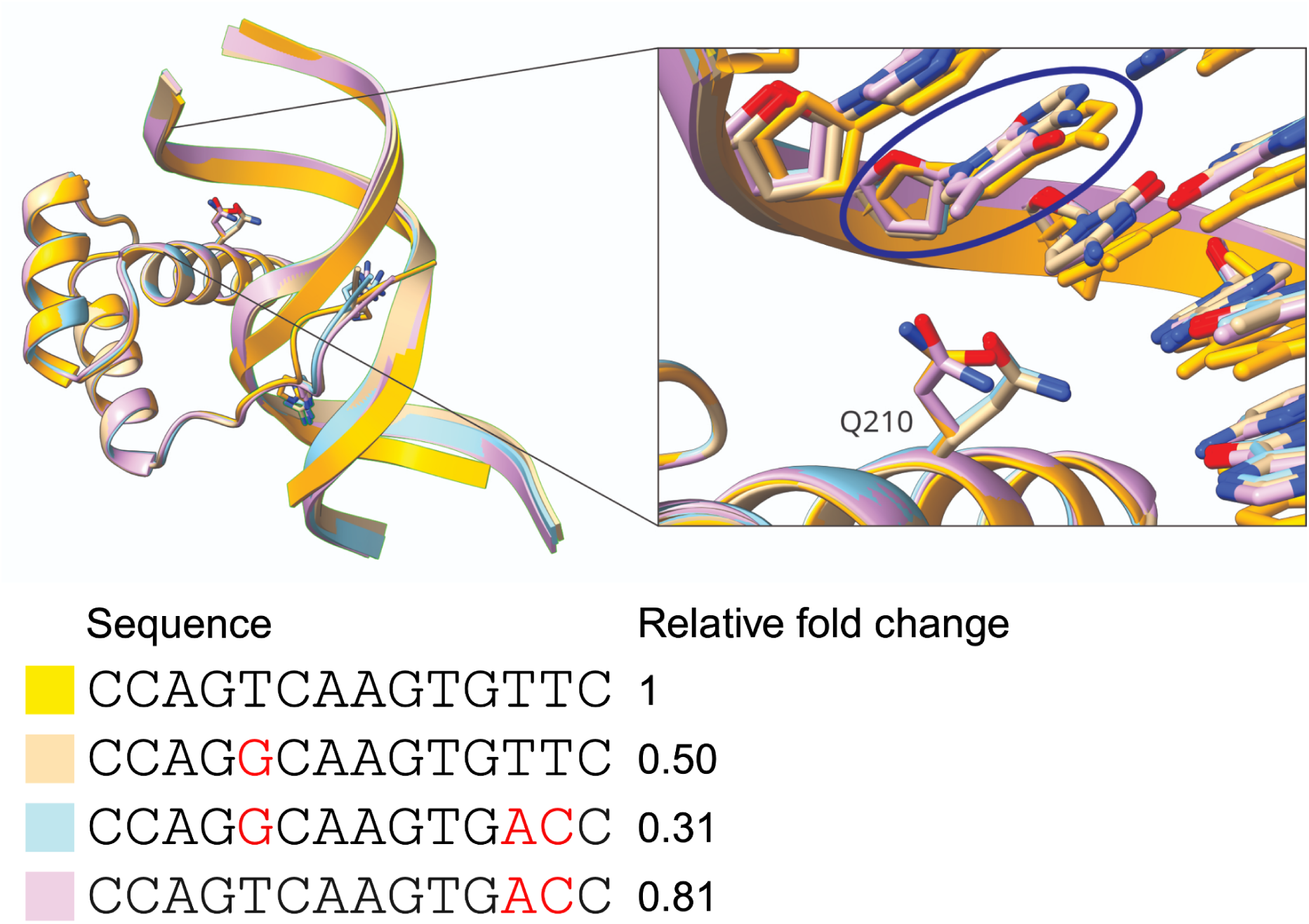
AlphaFold predictions for DNA sequences that had not been solved experimentally. The different NKX2.1 DBD-DNA sequence interactions are coloured according to the varied DNA sequences. The sequences are shown with their predicted *relative fold changes* by the **FLANK** model. The wildtype sequence structure is yellow, and the alternative bases for the other sequences are shown in red. The core sequence is boxed by a broken line.

### EMSA-seq derived models are able to identify NKX2.1 binding sites in genomic data

MST and selective chromatography [90] experiments provided a partial *in vitro* validation for our EMSA-seq-derived models. Next, we sought to test the models’ ability to identify NKX2.1 binding sites in genomic data. We conducted an *in vivo* validation experiment using publicly available, high-confidence ChIP-seq data for NKX2.1 and four negative control TFs (GATA1, MYOD1, NKX2.5, and RXRA). The peaks for NKX2.1 and the respective negative control TF were combined and the models used as classifiers to identify which TF each ChIP-seq peak belonged to, based on their LFC predictions. The peaks were filtered to remove any peaks that contained overlapping genomic regions to avoid situations where the two TFs might bind in the same region and skew results. We performed this classification task in two ways. First, we used the entire 500 bp sequence for our model LFC predictions (**Figure 9**), taking advantage of the models’ ability to take variable sequence lengths. Second, we used our models with a sliding window of 24 bp (the training sequence length) across the 500 bp ChIP-seq sequences and used the maximum scoring 24-mer to represent the peak (**Figure 10**). In both tests, the PWM was used with a 7 bp sliding window according to its length, and the maximum LFC was used to represent each peak. Despite not being trained for this task, our models performed relatively well (**Figures 9 and 10**). Unsurprisingly, the **CORE** model performed the worst. The restricted number of nucleotides varied in this training dataset restricts the length and complexity of patterns that can be learnt by the model. The **ALL** model performed better than the **CORE** model, and was mostly on par with the PWM, which was also not trained for this classification task. We expected this model to perform the best, owing to the high sequence diversity of the mutant library used in EMSA-seq. However, the high diversity of the **ALL** library had resulted in the requirement of a much higher read depth in order to achieve the same read-depth-to-unique-sequence ratio as compared to the other libraries, while the **FLANK** mutant library with its total variation of only 10 bps allowed detection of complex patterns at much lower total read depths. Hence, the **FLANK** model out-performed all other models, including the PWM, at the task of ChIP-seq peak classification. Interestingly, the performance of our models increased when using the entire 500 bp sequence (**Figure 9**) as compared to a 24 bp sliding window only (**Figure 10**). This is likely due to the increased sequence context included when using the entire peak instead of a sliding window, allowing the VCNNBPNet models to use nucleotide dependencies to make more accurate binding affinity predictions. The results from this classification task demonstrates that deep learning models trained on EMSA-seq data can predict binding *in vivo*.

**Figure 9:**
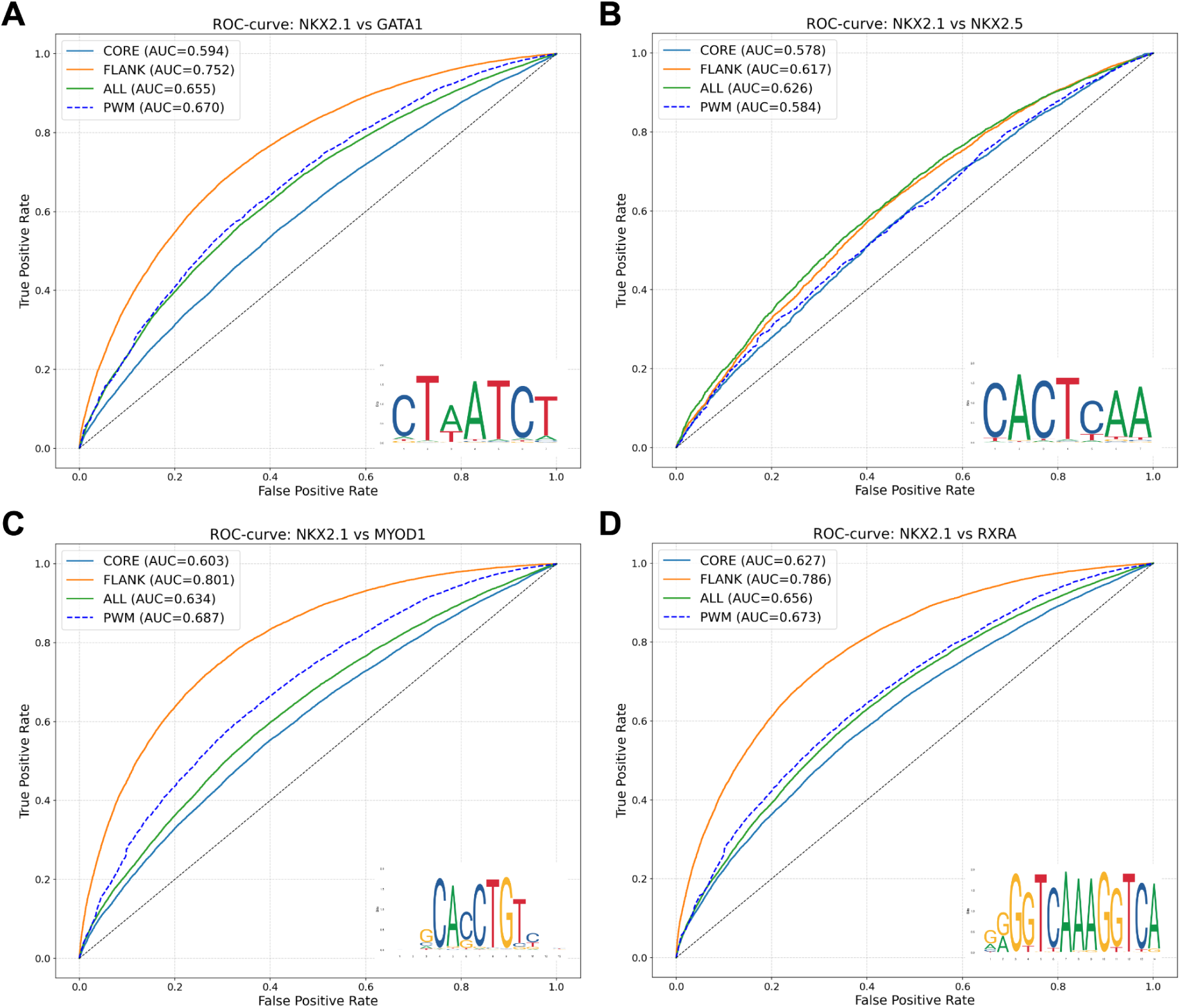
ChIP-seq peak classification by the PWM and VCNNBPNet models using a 500 bp DNA sequence. The classification task is performed using predicted LFCs for the ChIP-seq peak 500 bp sequences of NKX2.1 and negative control TFs **A:** GATA1, **B:** MYOD1, **C:** RXRA, and **D:** NKX2.5. The VCNNBPNet models were provided with the entire 500 bp ChIP-seq sequence as input and predicted the LFCs. Sequences of length 500 bp cannot be used as input for `FIMO` so the PWM (7 bp) was moved as a sliding window across the peak, and the highest scoring sequence was used as a representation for the entire peak. Receiver operator characteristic (ROC) curves are depicted to illustrate the models’ ability to distinguish between a positive (NKX2.1) and negative (other TF) dataset. The area under the curve (AUC) is stated for each model for each task. Each plot includes the negative control TF PWM from JASPAR [69] to visualise similarities between TFBSs.

**Figure 10:**
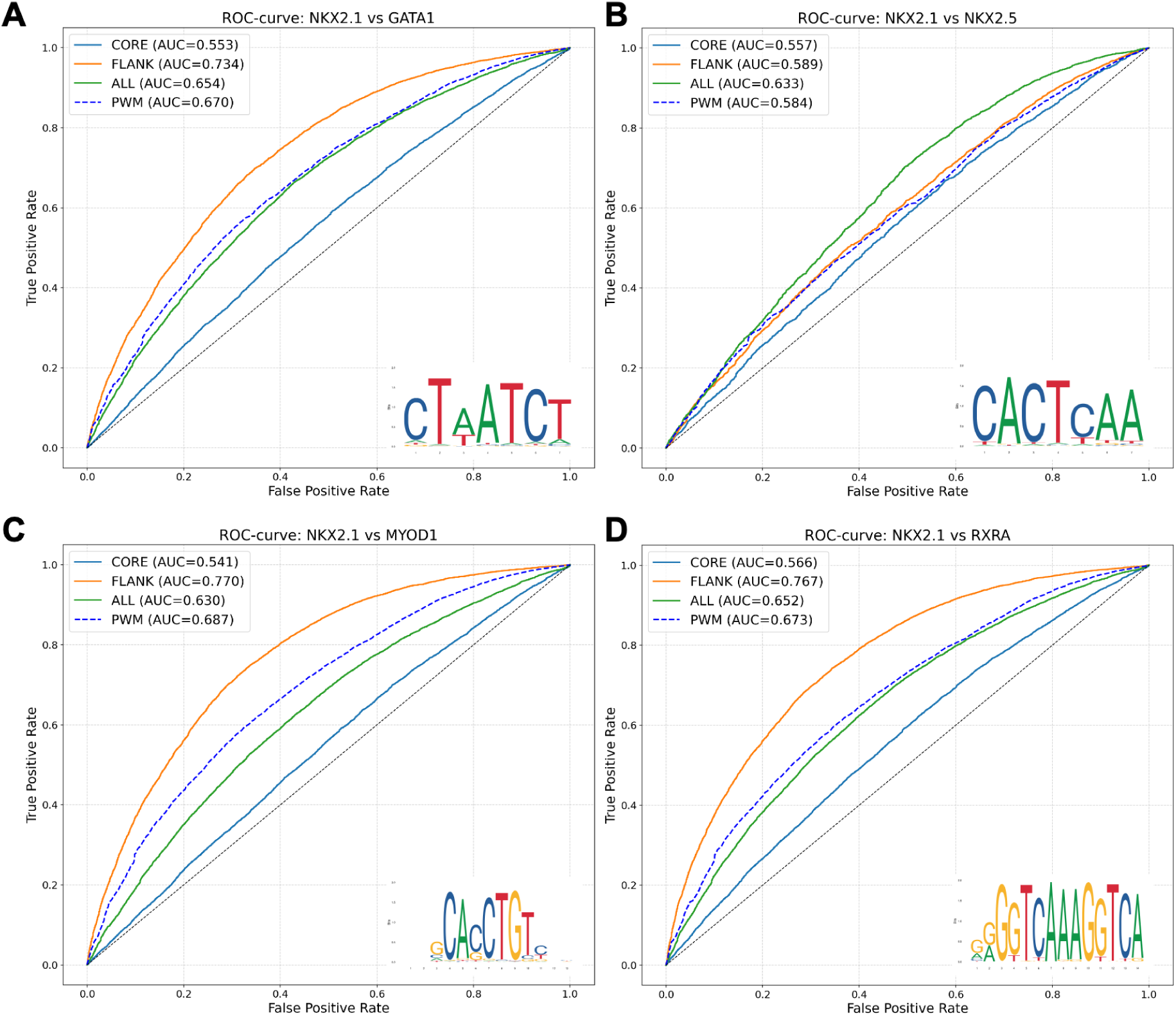
ChIP-seq peak classification by the PWM and VCNNBPNet models using a 24 bp sliding window. The classification task is performed using predicted LFCs for the ChIP-seq peak sequences of NKX2.1 and negative control TFs **A:** GATA1, **B:** MYOD1, **C:** RXRA, and **D:** NKX2.5. The VCNNBPNet models were provided with a 24 bp sliding window across the 500 bp peak sequence in 1 bp increments. The highest scoring 24 bp sequence was used as a representation of the peak. Sequences of 24 bp length cannot be used as input for `FIMO` so the PWM (7 bp) was moved as a sliding window across the peak, and the highest scoring sequence was used as a representation for the entire peak. Receiver operator characteristic (ROC) curves are depicted to illustrate the models’ ability to distinguish between a positive (NKX2.1) and negative control (other TF) dataset. The area under the curve (AUC) is stated for each model for each task. Each plot includes the negative control TF PWM from JASPAR [69] to visualise similarities between TFBSs.

**Figure 10:**
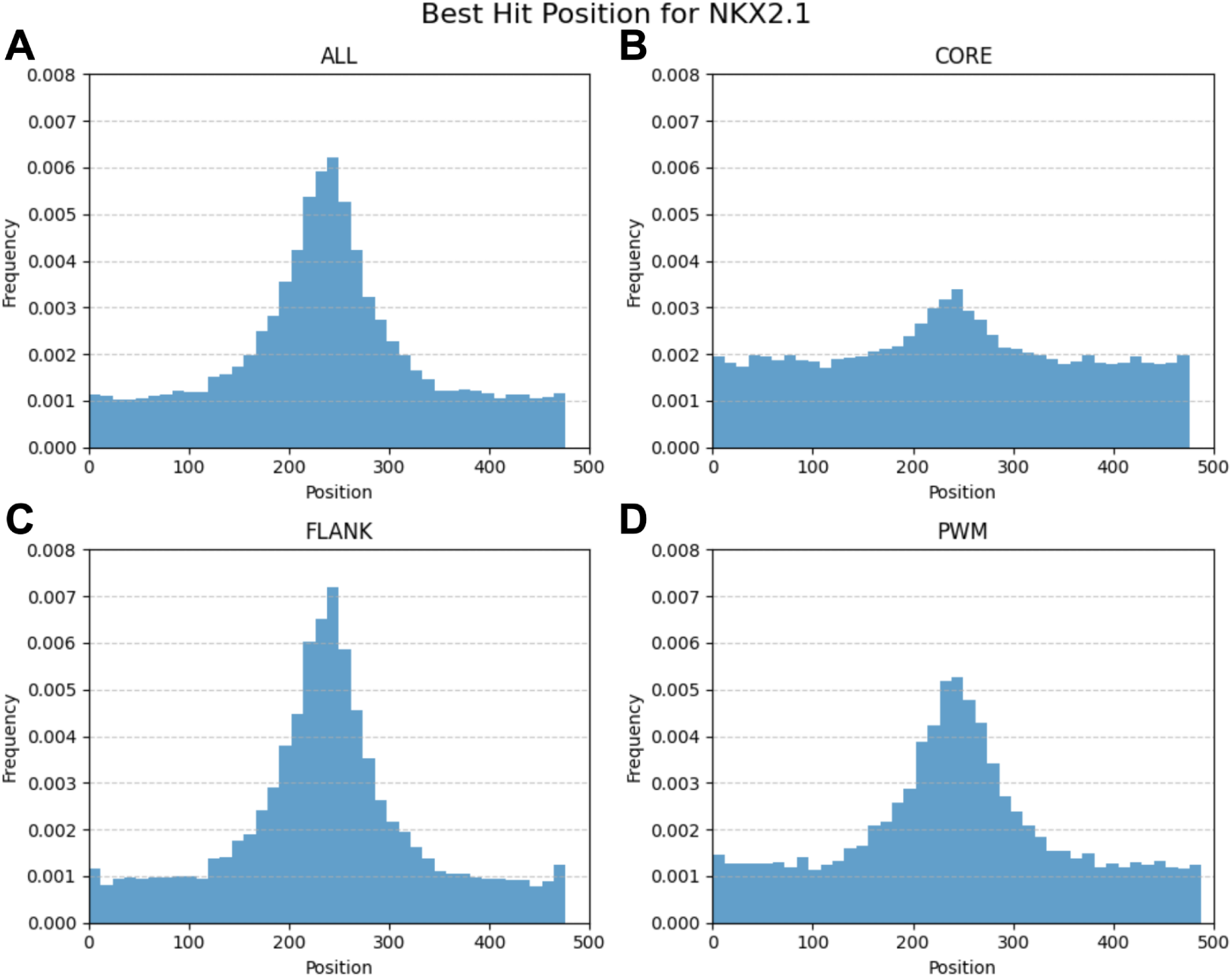
Highest model LFC prediction peak distribution plots for NKX2.1 ChIP-seq data. Each model was used to record where in the 500 bp NKX2.1 ChIP-seq peak sequence, the 24 bp DNA sequence with the highest predicted LFC occurred. The frequency of the highest predicted LFCs was plotted against the position this sequence was found in the peaks. **A: ALL** model, **B: CORE** model, **C: FLANK** model, and **D:** `FIMO` (length of sliding window was the size of the PWM).

In addition to the ChIP-seq peak classification task, we investigated where in the 500 bp NKX2.1 ChIP-peak sequence our models and the PWM model predicted the highest LFC (**Figure 10**). We hypothesised that well-performing models would have a high frequency of top LFCs in the centre of the peak sequence. For this task, the models assessed each 24-mer across the peak, in 1 bp increments, resulting in 477 positions. All models found the highest frequency of top LFCs in the centre of the NKX2.1 peaks. The **CORE** model (**Figure 10B**) performed the worst at this task, as for the ChIP-seq classification task. The **FLANK** (**Figure 10A**) and **ALL** (**Figure 10C**) models produced clear centre peaks, with the PWM’s (**Figure 10D**) centre peak slightly less pronounced. This task is based on the assumption that ChIP-seq peaks contain just one TFBS, and that the binding site should be at the centre of the peak, which is not always the case. Nevertheless, these results demonstrate the superiority of the **FLANK** model in NKX2.1 binding prediction *in vivo* compared to the PWM, despite a low correlation with MST binding affinity data.

### Final conclusions

We present the first publicly available, *in vitro* binding data between millions of DNA sequences and the human transcription factor NKX2.1. In addition, we trained neural network models that can accurately predict the relative binding strength of variable length DNA sequences, allowing prediction of the effects of nucleotide variants in known TFBSs. We also present direct binding affinity measurements between NKX2.1 and its thyroglobulin promoter binding site sequence and several introduced variants. Finally, we solved the crystal structure of the DNA-binding domain of NKX2.1 bound to its thyroglobulin promoter binding sequence and introduced several variants and used this to validate `AlphaFold` structure predictions for other DNA variants. This deep investigation of the binding between NKX2.1 and DNA revealed that *in vitro* EMSA-seq experiments can be used to train models with the ability of identifying TFBSs *in vivo* and that different methods for investigating TF binding can result in variable results (potentially due to the competitive nature of high-throughput methods). Future work should investigate the role of competition in binding reactions for other TFs. Importantly, this work allows the prediction of variants within known NKX2.1 binding sites, and could be used to assess variation in regulatory elements in patients with CAHTP in whom no mutation in the coding regions of the *NKX2.1* gene had been found.

## Acknowledgements

We would like to thank Raluca Gordân, PhD (Duke University School of Medicine) for her experimental discussion and advice.

## Author contribution

**Conceptualisation**: UH, MM, DS, MS; **Data curation:** SP, MS; **Formal analysis:** FLG, SP, MM; **Funding acquisition**: UH, DS, MS; **Investigation:** FLG, MM, MB; **Methodology:** FLG, MM, SP, MS, MB; **Project administration:** FLG, MS; **Resources:** UH, MS, DS; **Software:** SP; **Supervision:** MS, DS, UH; **Validation:** FLG, MM, SP; **Visualisation:** FLG, MM, SP; **Writing – original draft:** FLG, SP, MM; **Writing – review and editing:** all authors.

## Conflicts of interest

The authors do not report any conflict of interest with regard to this study.

## Data availability statement

Raw and processed data (*.`fastq` files, sequence counts, and `DESeq2` results) from the Illumina sequencing of the **CORE**, **FLANK**, and **ALL** libraries have been submitted to the Gene Expression Omnibus (GEO, https://www.ncbi.nlm.nih.gov/geo/) under the accession number GSE319141. The atomic coordinates and structure factors for the crystal structures described in this study have been deposited in the **Protein Data Bank (PDB)** under the following accession codes: **9U18** and **9U19.**

## Funding details

This research was funded by grants of the Deutsche Forschungsgemeinschaft (DFG) to MS within the NeuroCure Cluster of Excellence (EXC2049-390688087) and by DFG Research Unit FOR 2841 “Beyond the exome” (#400728090) to UH, HK, DS, and MS. This research was funded in part by the National Science Centre, Poland 3033/44/C/NZ1/00085 grant to MM.

## Notes

https://www.ncbi.nlm.nih.gov/geo/query/acc.cgi?acc=GSE319141

### Competing Interest Statement

The authors have declared no competing interest.

### Summary of Updates

Figures 5 and 8 have been corrected and updated.

