## Supplementary material for "Quantification of the effects of single nucleotide variants in NKX2.1 transcription factor binding sites"

### Supplementary materials

| # | Oligo name | Sequence [5'–3'] |
|---|---|---|
| **1** | Microscale thermophoresis dsDNA sequences | CACTGCCCA**XXXXXXXX**TTCTTGA |
| **2** | X-ray crystallography dsDNA sequences | AGT**XXXX**TGTTC |
| **3** | EMSA-seq **CORE** library | TCGTCGGCAGCGTCAGATGTGTATCACTGCCCAGT**NNNN**TGTTCTTGATCACCGCTCCGACTGCAGAAAA |
| **4** | EMSA-seq **FLANK** library | TCGTCGGCAGCGTCAGATGTGTATCACTGC**NNNNN**CAAG**NNNNN**TTGATCACCGCTCCGACTGCAGAAAA |
| **5** | EMSA-seq **ALL** library | TCGTCGGCAGCGTCAGATGTGTATCACTGC**NNNNNNNNNNNNNN**TTGATCACCGCTCCGACTGCAGAAAA |
| **6** | EMSA-seq positive control | TGCAGAACAGGTTTGTCTTTTGCTCACTGCCCAGTCAAGTGTTCTTGAGCAGAACTTAGAGGCGAGGTACGA |
| **7** | EMSA-seq negative control | CGTTACGACAGCAGAGTAGTGATGTGGCCTGGGCCCCTACGCGGCCGCCTGTAAGACTCACTCATCTCGGCT |
| **8** | EMSA-seq mutant library dsDNA preparation reverse primer | Alexa Fluor 647-TTTTCTGCAGTCGGAGCGGTGA |
| **9** | Negative control forward primer | CGTTACGACAGCAGAGTAGTGATG |
| **10** | Negative control reverse primer | AGCCGAGATGAGTGAGTCTTACAG |
| **11** | Positive control forward primer | ACTGTCTGAGGAGTGTCGTGTGCT |
| **12** | Positive control reverse primer | GTGAGTGAAGTGGTGCTGAGTCGT |
| **13** | Forward Illumina library preparation primer | AATGATACGGCGACCACCGAGATCTACACTAGATCGCTCGTCGGCAGCGTCAGATGTGTAT |
| **14** | Reverse Illumina library preparation primer (Ns indicate location of unique index) | CAAGCAGAAGACGGCATACGAGAT**NNNNNNNN**GTCTCGTGGGCTCGGAGATGTG |
| **15** | Illumina read primer 2 (used for **FLANK** and **ALL** libraries) | TCGTCGGCAGCGTCAGATGTGTATCACTGC |
| **16** | Illumina indexing primer | TCACCGCTCCGACTGCAGAAAA |

**Supplementary Table S1:** Oligonucleotides used in this study. N, positions where equal ratios of nucleotides (A:25, C:25, G:25, T:25) were incorporated randomly during DNA synthesis; X, specific nucleotide combinations were introduced (i.e. not every possible combination was used). Underlined nucleotides are primer sequences used for qPCR and Illumina sequencing library preparation.

| Category | Package Name | Version | Installation source |
|---|---|---|---|
| Bioinformatics | `bedtools` | 2.30.0 | System |
| Bioinformatics | `FIMO` | 5.5.7 | System |
| Bioinformatics (Python) | `biopython` | 1.85 | Pip |
| Bioinformatics (Python) | `logomaker` | 0.8 | Pip |
| Bioinformatics (Python) | `modisco-lite` | 2.2.1 | Pip |
| Bioinformatics (Python) | `pybigwig` | 0.3.23 | Pip |
| Bioinformatics (Python) | `pyfaidx` | 0.8.1.3 | Pip |
| Bioinformatics (Python) | `tangermeme` | 0.4.2 | Pip |
| Data Science & Viz | `altair` | 5.4.1 | Pip |
| Data Science & Viz | `bokeh` | 3.7.0 | Pip |
| Data Science & Viz | `h5py` | 3.12.1 | Pip |
| Data Science & Viz | `matplotlib` | 3.10.1 | Pip |
| Data Science & Viz | `networkx` | 3.1 | Pip |
| Data Science & Viz | `numpy` | 1.26.4 | Pip |

| | | | |
|---|---|---|---|
| Data Science & Viz | pandas | 2.2.2 | Pip |
| Data Science & Viz | plotly | 6.0.1 | Pip |
| Data Science & Viz | rich | 13.7.1 | Pip |
| Data Science & Viz | scipy | 1.14.1 | Conda |
| Data Science & Viz | seaborn | 0.13.2 | Pip |
| EMSA-seq data processing | apeglm | 1.32.0 | CRAN |
| EMSA-seq data processing | DESeq2 | 1.50.0 | Bioconductor |
| EMSA-seq data processing | ggplot2 | 4.0.0 | CRAN |
| EMSA-seq data processing | pheatmap | 1.0.13 | CRAN |
| EMSA-seq data processing | RColorBrewer | 1.1-3 | CRAN |
| EMSA-seq data processing | tidyverse | 2.0.0 | CRAN |
| Hardware (GPU/CUDA) | nvidia-cuda-runtime-cu12 | 12.1.105 | Pip |
| Hardware (GPU/CUDA) | nvidia-cudnn-cu12 | 9.1.0.70 | Pip |
| Jupyter Ecosystem | ipykernel | 6.15.0 | Conda |
| Jupyter Ecosystem | ipython | 8.14.0 | Conda |
| Jupyter Ecosystem | jupyterlab | 4.0.6 | Conda |
| Jupyter Ecosystem | nbconvert | 7.8.0 | Conda |
| Jupyter Ecosystem | notebook | 7.0.4 | Conda |
| Machine Learning & AI | lightning | 2.5.0 | Pip |
| Machine Learning & AI | pytorch-lightning | 2.5.0 | Pip |
| Machine Learning & AI | scikit-learn | 1.5.2 | Conda |
| Machine Learning & AI | shap | 0.48.0 | Pip |
| Machine Learning & AI | torch | 2.4.0 | Pip |
| Machine Learning & AI | torchmetrics | 1.4.1 | Pip |
| Machine Learning & AI | wandb | 0.17.6 | Pip |
| System & Core | pip | 23.1.2 | Conda |
| System & Core | python | 3.11.10 | Conda |
| System & Core | requests | 2.29.0 | Conda |
| System & Core | tqdm | 4.65.0 | Conda |
| System Libraries | gcc_impl_linux-64 | 14.2.0 | Conda |
| System Libraries | gstreamer | 1.24.7 | Conda |
| System Libraries | openssl | 3.4.1 | Conda |
| System Libraries | xorg-libx11 | 1.8.10 | Conda |

**Supplementary Table S2:** Software and software modules used in this project.

| Name | Created | Run time | Kernel _size | Model. kernel _size | Model. num_ channels | Model. pool _output _size | Num _channels | Pool _output _size | epoch | Train _loss _epoch | Train _loss _step | trainer/ global _step | val_loss |
|---|---|---|---|---|---|---|---|---|---|---|---|---|---|
| rare-sweep-16 | 2025-10-14T18:16:48.000Z | 44 | 13 | 13 | 82 | 178 | 82 | 178 | 36 | 0.092099555 | 0.052138545 | 221 | 0.06079312 |
| generous-sweep-76 | 2025-10-14T19:05:21.000Z | 45 | 36 | 36 | 80 | 209 | 80 | 209 | 40 | 0.050105657 | 0.039100494 | 245 | 0.06141796 |
| earnest-sweep-40 | 2025-10-14T18:35:36.000Z | 43 | 15 | 15 | 77 | 92 | 77 | 92 | 43 | 0.027660601 | 0.034931377 | 263 | 0.06260511 |
| dainty-sweep-96 | 2025-10-14T19:21:08.000Z | 41 | 28 | 28 | 74 | 84 | 74 | 84 | 38 | 0.055511702 | 0.023969289 | 233 | 0.07203593 |
| lunar-sweep-4 | 2025-10-14T18:07:18.000Z | 43 | 12 | 12 | 98 | 108 | 98 | 108 | 45 | 0.03395161 | 0.03194388 | 275 | 0.07892106 |
| sleek-sweep-56 | 2025-10-14T18:49:21.000Z | 40 | 13 | 13 | 97 | 94 | 97 | 94 | 33 | 0.035093531 | 0.027831677 | 203 | 0.08082398 |
| feasible-sweep-66 | 2025-10-14T18:57:09.000Z | 42 | 14 | 14 | 104 | 173 | 104 | 173 | 45 | 0.041814271 | 0.025270987 | 275 | 0.08877626 |
| clear-sweep-52 | 2025-10-14T18:45:48.000Z | 41 | 17 | 17 | 74 | 214 | 74 | 214 | 50 | 0.035827693 | 0.015849298 | 305 | 0.09375614 |
| bumbling-sweep-25 | 2025-10-14T18:23:55.000Z | 42 | 23 | 23 | 85 | 73 | 85 | 73 | 43 | 0.058448948 | 0.07701762 | 263 | 0.0940453 |
| icy-sweep-19 | 2025-10-14T18:19:14.000Z | 39 | 22 | 22 | 72 | 65 | 72 | 65 | 35 | 0.095430799 | 0.033007555 | 215 | 0.09645522 |
| eager-sweep-22 | 2025-10-14T18:21:34.000Z | 41 | 21 | 21 | 46 | 79 | 46 | 79 | 46 | 0.074601941 | 0.124091104 | 281 | 0.09753805 |
| ancient-sweep-30 | 2025-10-14T18:27:49.000Z | 42 | 33 | 33 | 42 | 187 | 42 | 187 | 29 | 0.064417042 | 0.138720125 | 179 | 0.09792756 |
| fallen-sweep-41 | 2025-10-14T18:36:23.000Z | 43 | 24 | 24 | 84 | 71 | 84 | 71 | 39 | 0.044630457 | 0.014613479 | 239 | 0.09877452 |
| eager-sweep-1 | 2025-10-14T18:04:37.000Z | 57 | 14 | 14 | 93 | 196 | 93 | 196 | 32 | 0.103157468 | 0.077240862 | 197 | 0.10162681 |
| vocal-sweep-5 | 2025-10-14T18:08:05.000Z | 41 | 14 | 14 | 106 | 139 | 106 | 139 | 25 | 0.092669308 | 0.059204038 | 155 | 0.10633692 |
| resilient-sweep-2 | 2025-10-14T18:05:39.000Z | 47 | 13 | 13 | 85 | 89 | 85 | 89 | 39 | 0.069771461 | 0.067813501 | 239 | 0.10813407 |
| woven-sweep-7 | 2025-10-14T18:09:39.000Z | 42 | 23 | 23 | 41 | 84 | 41 | 84 | 40 | 0.068083309 | 0.042314664 | 245 | 0.11088385 |
| avid-sweep-46 | 2025-10-14T18:40:53.000Z | 43 | 32 | 32 | 88 | 221 | 88 | 221 | 31 | 0.060312331 | 0.020770879 | 191 | 0.11202669 |
| genial-sweep-84 | 2025-10-14T19:11:45.000Z | 39 | 17 | 17 | 51 | 157 | 51 | 157 | 35 | 0.078713223 | 0.046864759 | 215 | 0.11324269 |
| giddy-sweep-82 | 2025-10-14T19:10:01.000Z | 45 | 22 | 22 | 119 | 223 | 119 | 223 | 30 | 0.103911616 | 0.057643339 | 185 | 0.12130735 |
| vocal-sweep-36 | 2025-10-14T18:32:24.000Z | 45 | 41 | 41 | 105 | 244 | 105 | 244 | 38 | 0.037935 | 0.046485208 | 233 | 0.12511574 |

| | | | | | | | | | | | | | |
|---|---|---|---|---|---|---|---|---|---|---|---|---|---|
| dashing-sweep-17 | 2025-10-14T18:17:40.000Z | 42 | 12 | 12 | 84 | 214 | 84 | 214 | 37 | 0.121385433 | 0.155370533 | 227 | 0.1261221 |
| splendid-sweep-12 | 2025-10-14T18:13:33.000Z | 41 | 18 | 18 | 80 | 83 | 80 | 83 | 27 | 0.102476716 | 0.093761712 | 167 | 0.14570934 |
| fallen-sweep-68 | 2025-10-14T18:58:42.000Z | 41 | 17 | 17 | 89 | 229 | 89 | 229 | 37 | 0.077765495 | 0.065464415 | 227 | 0.14893766 |
| valiant-sweep-59 | 2025-10-14T18:51:41.000Z | 44 | 10 | 10 | 100 | 74 | 100 | 74 | 26 | 0.097072981 | 0.083956286 | 161 | 0.15515105 |
| sleek-sweep-86 | 2025-10-14T19:13:19.000Z | 38 | 32 | 32 | 40 | 127 | 40 | 127 | 25 | 0.205783024 | 0.270869613 | 155 | 0.16100238 |
| silver-sweep-11 | 2025-10-14T18:12:41.000Z | 44 | 15 | 15 | 79 | 170 | 79 | 170 | 33 | 0.226546347 | 0.501060903 | 203 | 0.16479965 |
| pleasant-sweep-88 | 2025-10-14T19:15:04.000Z | 38 | 11 | 11 | 45 | 255 | 45 | 255 | 22 | 0.169882029 | 0.134999529 | 137 | 0.16814552 |
| worthy-sweep-50 | 2025-10-14T18:44:00.000Z | 41 | 13 | 13 | 90 | 122 | 90 | 122 | 20 | 0.14156507 | 0.085205883 | 125 | 0.16936144 |
| dry-sweep-70 | 2025-10-14T19:00:21.000Z | 42 | 29 | 29 | 106 | 212 | 106 | 212 | 27 | 0.183797911 | 0.106086105 | 167 | 0.17228262 |
| major-sweep-77 | 2025-10-14T19:06:13.000Z | 40 | 10 | 10 | 41 | 83 | 41 | 83 | 24 | 0.258832783 | 0.328919888 | 149 | 0.17457601 |
| earthy-sweep-8 | 2025-10-14T18:10:25.000Z | 43 | 19 | 19 | 98 | 83 | 98 | 83 | 19 | 0.248645246 | 0.528274536 | 119 | 0.17834279 |
| generous-sweep-62 | 2025-10-14T18:54:07.000Z | 41 | 31 | 31 | 78 | 153 | 78 | 153 | 35 | 0.088540852 | 0.04771265 | 215 | 0.18333368 |
| dark-sweep-23 | 2025-10-14T18:22:21.000Z | 43 | 11 | 11 | 83 | 176 | 83 | 176 | 29 | 0.464718968 | 0.059250154 | 179 | 0.19072567 |
| glowing-sweep-6 | 2025-10-14T18:08:52.000Z | 42 | 10 | 10 | 86 | 117 | 86 | 117 | 23 | 0.10110683 | 0.096215166 | 143 | 0.193546 |
| true-sweep-3 | 2025-10-14T18:06:31.000Z | 43 | 38 | 38 | 87 | 211 | 87 | 211 | 22 | 0.201571524 | 0.266815126 | 137 | 0.19723153 |
| astral-sweep-35 | 2025-10-14T18:31:37.000Z | 41 | 10 | 10 | 74 | 69 | 74 | 69 | 26 | 0.159597248 | 0.107509956 | 161 | 0.19942515 |
| comic-sweep-44 | 2025-10-14T18:38:49.000Z | 47 | 17 | 17 | 118 | 178 | 118 | 178 | 42 | 0.09949059 | 0.138869986 | 257 | 0.20102094 |
| clean-sweep-97 | 2025-10-14T19:21:55.000Z | 43 | 17 | 17 | 128 | 237 | 128 | 237 | 22 | 0.201687798 | 0.214614123 | 137 | 0.20114811 |
| deep-sweep-61 | 2025-10-14T18:53:20.000Z | 42 | 11 | 11 | 51 | 99 | 51 | 99 | 22 | 0.175707683 | 0.167355657 | 137 | 0.21037772 |
| celestial-sweep-31 | 2025-10-14T18:28:35.000Z | 41 | 16 | 16 | 65 | 148 | 65 | 148 | 20 | 0.210957915 | 0.130622759 | 125 | 0.21257144 |
| zesty-sweep-65 | 2025-10-14T18:56:22.000Z | 41 | 27 | 27 | 122 | 98 | 122 | 98 | 20 | 0.172561899 | 0.049167894 | 125 | 0.22467786 |
| fiery-sweep-93 | 2025-10-14T19:18:53.000Z | 38 | 31 | 31 | 72 | 75 | 72 | 75 | 20 | 0.312247545 | 0.075045511 | 125 | 0.22656938 |
| lilac-sweep-32 | 2025-10-14T18:29:22.000Z | 43 | 33 | 33 | 115 | 170 | 115 | 170 | 17 | 0.212438926 | 0.145025536 | 107 | 0.22749431 |
| vivid-sweep-87 | 2025-10-14T19:14:17.000Z | 42 | 17 | 17 | 89 | 81 | 89 | 81 | 30 | 0.110402443 | 0.053915456 | 185 | 0.23991743 |
| sweet-sweep-45 | 2025-10-14T18:39:41.000Z | 43 | 33 | 33 | 92 | 204 | 92 | 204 | 30 | 0.278908879 | 0.152278692 | 185 | 0.25460652 |

| | | | | | | | | | | | | | |
|---|---|---|---|---|---|---|---|---|---|---|---|---|---|
| fancy-sweep-20 | 2025-10-14T18:20:01.000Z | 41 | 14 | 14 | 75 | 71 | 75 | 71 | 24 | 0.136343464 | 0.03814042 | 149 | 0.25938007 |
| happy-sweep-26 | 2025-10-14T18:24:42.000Z | 41 | 14 | 14 | 80 | 205 | 80 | 205 | 17 | 0.207686484 | 0.194484442 | 107 | 0.27681306 |
| still-sweep-57 | 2025-10-14T18:50:08.000Z | 42 | 12 | 12 | 93 | 146 | 93 | 146 | 16 | 0.241158143 | 0.140067652 | 101 | 0.27777728 |
| polar-sweep-42 | 2025-10-14T18:37:10.000Z | 43 | 15 | 15 | 73 | 108 | 73 | 108 | 30 | 0.38114816 | 0.09340331 | 185 | 0.28933746 |
| vibrant-sweep-69 | 2025-10-14T18:59:29.000Z | 43 | 30 | 30 | 101 | 156 | 101 | 156 | 19 | 0.20735459 | 0.164394647 | 119 | 0.31023651 |
| playful-sweep-24 | 2025-10-14T18:23:08.000Z | 39 | 16 | 16 | 43 | 127 | 43 | 127 | 29 | 0.411148876 | 0.027755052 | 179 | 0.32302809 |
| solar-sweep-78 | 2025-10-14T19:06:59.000Z | 38 | 27 | 27 | 38 | 154 | 38 | 154 | 17 | 0.373859853 | 0.345107615 | 107 | 0.33810845 |
| floral-sweep-58 | 2025-10-14T18:50:55.000Z | 42 | 15 | 15 | 100 | 182 | 100 | 182 | 27 | 0.171617448 | 0.509374619 | 167 | 0.35908693 |
| legendary-sweep-27 | 2025-10-14T18:25:28.000Z | 42 | 12 | 12 | 92 | 97 | 92 | 97 | 17 | 0.237132609 | 0.311489999 | 107 | 0.37864071 |
| pleasant-sweep-13 | 2025-10-14T18:14:28.000Z | 41 | 19 | 19 | 90 | 157 | 90 | 157 | 20 | 0.558061719 | 0.070564911 | 125 | 0.38354969 |
| rural-sweep-55 | 2025-10-14T18:48:29.000Z | 45 | 29 | 29 | 126 | 105 | 126 | 105 | 25 | 0.417155832 | 0.076968536 | 155 | 0.40800485 |
| super-sweep-18 | 2025-10-14T18:18:27.000Z | 42 | 17 | 17 | 89 | 118 | 89 | 118 | 21 | 0.373695791 | 0.573626935 | 131 | 0.4227708 |
| worldly-sweep-54 | 2025-10-14T18:47:22.000Z | 40 | 34 | 34 | 39 | 85 | 39 | 85 | 17 | 0.335975021 | 0.265072048 | 107 | 0.42611802 |
| playful-sweep-90 | 2025-10-14T19:16:32.000Z | 38 | 36 | 36 | 89 | 145 | 89 | 145 | 13 | 0.546335816 | 0.387940019 | 83 | 0.47274747 |
| earthy-sweep-10 | 2025-10-14T18:11:59.000Z | 37 | 11 | 11 | 117 | 113 | 117 | 113 | 13 | 0.460335344 | 1.519062757 | 83 | 0.48649496 |
| usual-sweep-60 | 2025-10-14T18:52:33.000Z | 42 | 17 | 17 | 123 | 248 | 123 | 248 | 22 | 0.369553268 | 0.249619186 | 137 | 0.51050031 |
| electric-sweep-94 | 2025-10-14T19:19:39.000Z | 38 | 21 | 21 | 82 | 232 | 82 | 232 | 26 | 0.306021899 | 0.336771607 | 161 | 0.51620746 |
| eternal-sweep-43 | 2025-10-14T18:38:02.000Z | 39 | 19 | 19 | 88 | 79 | 88 | 79 | 21 | 0.561744332 | 1.389275432 | 131 | 0.52278948 |
| fine-sweep-49 | 2025-10-14T18:43:13.000Z | 39 | 24 | 24 | 69 | 214 | 69 | 214 | 22 | 0.551522255 | 1.655740499 | 137 | 0.52992803 |
| elated-sweep-75 | 2025-10-14T19:04:34.000Z | 40 | 17 | 17 | 128 | 101 | 128 | 101 | 20 | 0.486647308 | 0.38857162 | 125 | 0.5370307 |
| earthy-sweep-47 | 2025-10-14T18:41:40.000Z | 38 | 15 | 15 | 111 | 184 | 111 | 184 | 18 | 0.592418075 | 0.20092541 | 113 | 0.54657716 |
| vital-sweep-98 | 2025-10-14T19:22:42.000Z | 38 | 26 | 26 | 102 | 119 | 102 | 119 | 24 | 0.703357279 | 0.248088241 | 149 | 0.56440133 |
| pleasant-sweep-99 | 2025-10-14T19:23:29.000Z | 38 | 29 | 29 | 43 | 133 | 43 | 133 | 20 | 0.547877192 | 0.087132648 | 125 | 0.56682849 |
| curious-sweep-38 | 2025-10-14T18:34:03.000Z | 39 | 29 | 29 | 112 | 247 | 112 | 247 | 11 | 0.458060086 | 0.415527821 | 71 | 0.58450752 |
| comic-sweep-100 | 2025-10-14T19:24:15.000Z | 40 | 22 | 22 | 63 | 138 | 63 | 138 | 20 | 0.680905223 | 0.222422048 | 125 | 0.60747534 |

| | | | | | | | | | | | | | |
|---|---|---|---|---|---|---|---|---|---|---|---|---|---|
| fine-sweep-83 | 2025-10-14T19:10:53.000Z | 44 | 37 | 37 | 124 | 88 | 124 | 88 | 21 | 0.726376832 | 0.319216877 | 131 | 0.6378991 |
| summer-sweep-51 | 2025-10-14T18:45:01.000Z | 41 | 31 | 31 | 59 | 233 | 59 | 233 | 20 | 0.723827243 | 0.336398929 | 125 | 0.64508468 |
| sunny-sweep-85 | 2025-10-14T19:12:32.000Z | 41 | 37 | 37 | 35 | 189 | 35 | 189 | 19 | 0.722772181 | 0.295447856 | 119 | 0.66011441 |
| genial-sweep-29 | 2025-10-14T18:27:02.000Z | 38 | 27 | 27 | 47 | 88 | 47 | 88 | 12 | 0.543634593 | 0.479541332 | 77 | 0.68855208 |
| still-sweep-63 | 2025-10-14T18:54:54.000Z | 42 | 22 | 22 | 65 | 139 | 65 | 139 | 13 | 0.663317323 | 0.919211388 | 83 | 0.70890892 |
| still-sweep-71 | 2025-10-14T19:01:08.000Z | 43 | 13 | 13 | 91 | 84 | 91 | 84 | 24 | 0.80978626 | 0.739283979 | 149 | 0.72018272 |
| floral-sweep-33 | 2025-10-14T18:30:09.000Z | 41 | 37 | 37 | 35 | 215 | 35 | 215 | 13 | 0.81628406 | 0.973987937 | 83 | 0.7256211 |
| still-sweep-37 | 2025-10-14T18:33:16.000Z | 40 | 26 | 26 | 118 | 209 | 118 | 209 | 15 | 0.990578592 | 0.079417422 | 95 | 0.87831563 |
| glad-sweep-72 | 2025-10-14T19:02:00.000Z | 38 | 33 | 33 | 68 | 116 | 68 | 116 | 15 | 0.996387124 | 0.258243501 | 95 | 0.87917978 |
| sandy-sweep-34 | 2025-10-14T18:30:56.000Z | 37 | 26 | 26 | 117 | 98 | 117 | 98 | 5 | 0.999137282 | 0.103736036 | 35 | 0.88300771 |
| glowing-sweep-91 | 2025-10-14T19:17:14.000Z | 46 | 39 | 39 | 119 | 253 | 119 | 253 | 19 | 0.998232484 | 0.136228129 | 119 | 0.88309598 |
| vocal-sweep-21 | 2025-10-14T18:20:48.000Z | 39 | 29 | 29 | 44 | 78 | 44 | 78 | 18 | 0.984303832 | 0.631527662 | 113 | 0.8832258 |
| eager-sweep-79 | 2025-10-14T19:07:46.000Z | 41 | 22 | 22 | 103 | 124 | 103 | 124 | 13 | 0.997176349 | 1.133627057 | 83 | 0.88361543 |
| vague-sweep-67 | 2025-10-14T18:57:56.000Z | 40 | 30 | 30 | 116 | 255 | 116 | 255 | 5 | 0.996010721 | 5.174656868 | 35 | 0.88390696 |
| sandy-sweep-39 | 2025-10-14T18:34:49.000Z | 39 | 35 | 35 | 55 | 225 | 55 | 225 | 6 | 0.992671847 | 0.813077211 | 41 | 0.88474238 |
| stoic-sweep-89 | 2025-10-14T19:15:45.000Z | 39 | 13 | 13 | 58 | 165 | 58 | 165 | 7 | 0.9969064 | 0.491814911 | 47 | 0.88565797 |
| rosy-sweep-28 | 2025-10-14T18:26:15.000Z | 38 | 28 | 28 | 33 | 144 | 33 | 144 | 5 | 0.993139088 | 0.187875673 | 35 | 0.88584447 |
| vibrant-sweep-53 | 2025-10-14T18:46:35.000Z | 40 | 19 | 19 | 109 | 183 | 109 | 183 | 6 | 1.002271175 | 0.367721677 | 41 | 0.88670015 |
| kind-sweep-48 | 2025-10-14T18:42:26.000Z | 43 | 30 | 30 | 45 | 222 | 45 | 222 | 19 | 0.997078717 | 0.104206286 | 119 | 0.8876754 |
| dutiful-sweep-14 | 2025-10-14T18:15:14.000Z | 40 | 14 | 14 | 118 | 189 | 118 | 189 | 7 | 0.997144938 | 0.498690814 | 47 | 0.88827848 |
| brisk-sweep-74 | 2025-10-14T19:03:52.000Z | 36 | 11 | 11 | 63 | 229 | 63 | 229 | 5 | 0.997130811 | 0.248213708 | 35 | 0.88859791 |
| crimson-sweep-9 | 2025-10-14T18:11:12.000Z | 38 | 40 | 40 | 59 | 189 | 59 | 189 | 14 | 0.997266531 | 1.4479599 | 89 | 0.8888225 |
| eager-sweep-92 | 2025-10-14T19:18:06.000Z | 39 | 35 | 35 | 117 | 163 | 117 | 163 | 11 | 0.997123599 | 1.325618029 | 71 | 0.88888121 |
| eternal-sweep-81 | 2025-10-14T19:09:14.000Z | 41 | 32 | 32 | 51 | 137 | 51 | 137 | 5 | 0.998182774 | 0.149682894 | 35 | 0.88926977 |
| denim-sweep-64 | 2025-10-14T18:55:40.000Z | 36 | 34 | 34 | 63 | 244 | 63 | 244 | 6 | 0.997027516 | 1.010430336 | 41 | 0.88979542 |

| | | | | | | | | | | | | | |
|---|---|---|---|---|---|---|---|---|---|---|---|---|---|
| mild-sweep-73 | 2025-10-14T19:03:10.000Z | 38 | 11 | 11 | 96 | 88 | 96 | 88 | 5 | 0.99800843 | 0.1778934 | 35 | 0.89094591 |
| solar-sweep-15 | 2025-10-14T18:16:01.000Z | 40 | 36 | 36 | 43 | 242 | 43 | 242 | 17 | 0.998658538 | 1.338091612 | 107 | 0.89097923 |
| faithful-sweep-80 | 2025-10-14T19:08:33.000Z | 35 | 33 | 33 | 64 | 175 | 64 | 175 | 5 | 1.00009954 | 0.30474934 | 35 | 0.89459276 |
| blooming-sweep-95 | 2025-10-14T19:20:21.000Z | 42 | 15 | 15 | 127 | 247 | 127 | 247 | 9 | 1.001496792 | 0.18563664 | 59 | |

**Supplementary Table S3: Optimal hyperparameter settings for the VCNNBPNet models.** Note: these settings were identical for all models: `batch_size = 64`, `classify = FALSE`, `dense_sizes = [128,64]`, `dilations = [1,1,2,4,8]`, `input_channels = 4`, `input_length = 24`, `learning_rate = 0.001`, `model.dense_sizes = [128,64]`, `model.dilations = [1,1,2,4,8]`, `num_workers = 24`, `preprocess = FALSE`, and `scaling_method = standardise`.

| ChipSeq Target | GSM Accession IDs |
|---|---|
| **GATA1** | GSM1003608, GSM1067274, GSM1278240, GSM1816080, GSM1816081, GSM1921310, GSM1921315, GSM1921321, GSM1921327, GSM1921332, GSM2452102, GSM2877105, GSM2877106, GSM2877113, GSM2877114, GSM2877121, GSM2877122, GSM3177416, GSM3177417, GSM3177418, GSM3177419, GSM3177420, GSM3177421, GSM3177422, GSM3177423, GSM3523237, GSM3523244, GSM3762809, GSM3762810, GSM3762813, GSM3762817, GSM3783517, GSM3832646, GSM3832647, GSM3832653, GSM3832654, GSM3854067, GSM3854068, GSM3854083, GSM3854084, GSM3854099, GSM3854100, GSM3854115, GSM3854116, GSM3854131, GSM3854132, GSM4096234, GSM4096235, GSM4096236, GSM4613166, GSM4613167, GSM4613168, GSM4613169, GSM4613171, GSM4613172, GSM4613173, GSM4613174, GSM4613175, GSM4613176, GSM4613183, GSM4613184, GSM4613185, GSM4613186, GSM467647, GSM4761236, GSM4761237, GSM4818700, GSM4818708, GSM5129496, GSM5129497, GSM5129498, GSM5129499, GSM5129500, GSM5129501, GSM5271205, GSM5271206, GSM5271209, GSM5271210, GSM5271219, GSM5271220, GSM5271222, GSM5271223, GSM5341623, GSM5575500, GSM5575501, GSM5696922, GSM5696923, GSM5696924, GSM5696926, GSM5696927, GSM5696928, GSM5696930, GSM5696931, GSM5696932, GSM5771686, GSM5771687, GSM5771688, GSM5771689, GSM607949, GSM610335, GSM6321450, GSM6321451, GSM6321454, GSM6321455, GSM6463411, GSM6463412, GSM6463413, GSM651546, GSM651547, GSM6661430, GSM6661431, GSM6661432, GSM6661433, GSM722392, GSM722393, GSM722394, GSM722395, GSM722412, GSM722413, GSM788367, GSM804013, GSM804014, GSM935333, GSM935465, GSM935540, GSM970257, GSM970258 |
| **MYOD1** | GSM1218849, GSM1218850, GSM1218851, GSM1239474, GSM2214114, GSM2259152, GSM2259153, GSM2259154, GSM3831286, GSM3831287, GSM4072349, GSM4072350, GSM4072354, GSM4072355, GSM5261170, GSM5261171, GSM5261172, GSM6280737, GSM6280738, GSM6280739, GSM6280740, GSM6626408, GSM6626409, GSM6626410, GSM6626411, GSM6626416, GSM6626417, GSM6626418, GSM6626419, GSM7720563, GSM7720564 |
| **NKX2.1** | GSM1246715, GSM1246716, GSM2310996, GSM2310998, GSM5556351, GSM5556352, GSM5556353, GSM5556354, GSM5556355, GSM5556356, GSM5556357, GSM5556368, GSM5556369, GSM5556370, GSM5556371, GSM5556372, GSM5556373, GSM983098,<br>GSM983100, GSM983102, GSM983104 |
| **NKX2.5** | GSM2372592, GSM2372593, GSM2372594, GSM4829313, GSM4829316, GSM5838060, GSM5838061, GSM5838062, GSM5838077, GSM5838079 |
| **RXRA** | GSM1010767, GSM1019136, GSM1239520, GSM1480741, GSM2877302, GSM2877303, GSM2877304, GSM2877305, GSM2877306, GSM2877307, GSM3384456, GSM3384457, GSM3384458, GSM3384471, GSM3384472, GSM3384473, GSM3634227, GSM3634228, GSM3636221, GSM3636222, GSM4083801, GSM468181, GSM468184, GSM468199, GSM468204, GSM468213, GSM468214, GSM5085851, GSM5085852, GSM5213954, GSM5213955, GSM624142, GSM791405, GSM791406, GSM803341, GSM803452, GSM803506 |

**Supplementary Table S4:** Gene Expression Omnibus (https://www.ncbi.nlm.nih.gov/geo/) repository IDs of ChIP-seq data used for validation. The data was downloaded from ChIP-Atlas ([95]).

### Supplementary results

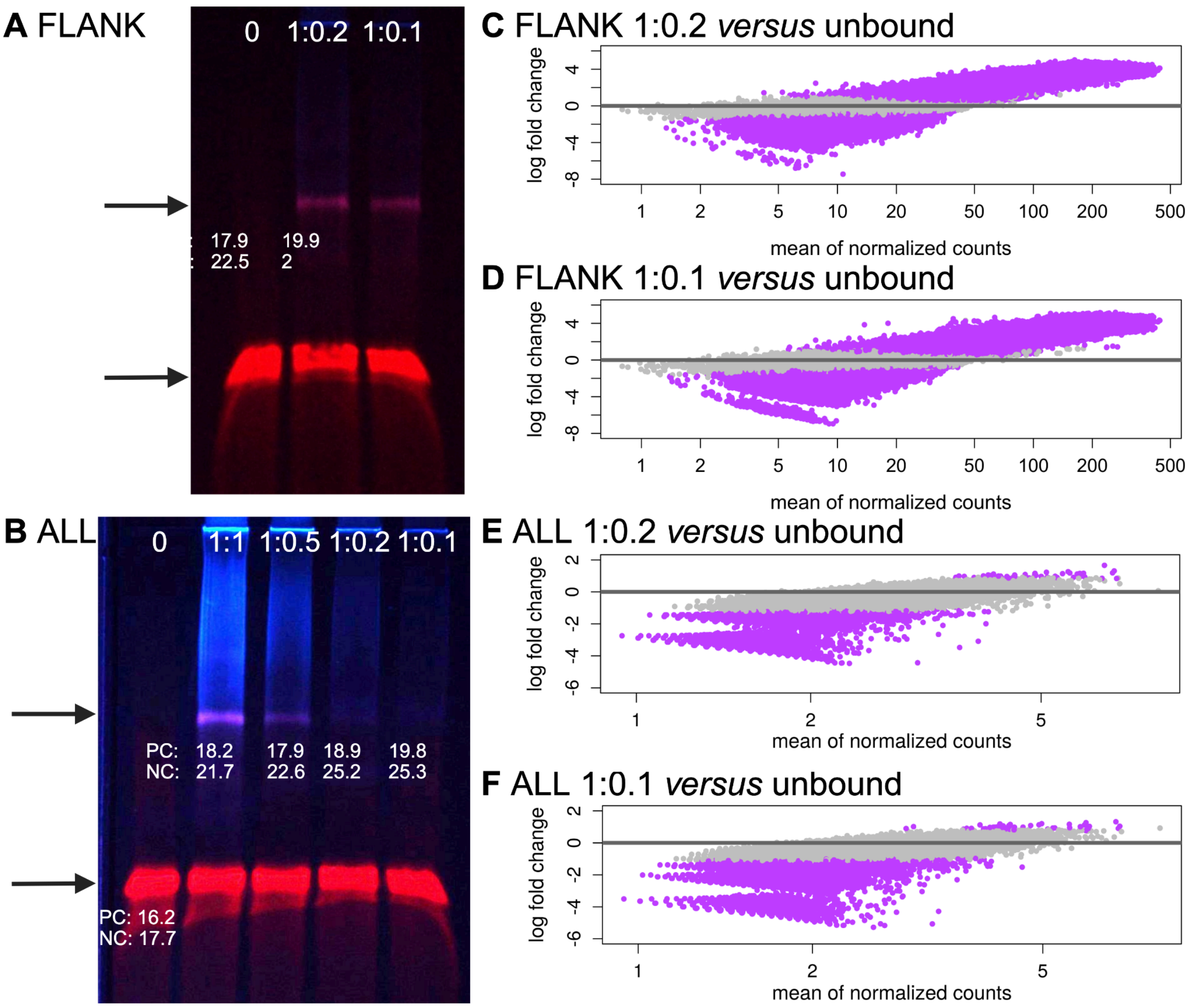


**Supplementary Figure S1:** Electromobility shift assays with NKX2.1 (blue) and **A: FLANK** (red) or **B: ALL** (red) mutant libraries. The DNA to protein molar ratios of each lane are indicated at the top (0 has no protein). The top arrow indicates NKX2.1-bound DNA and the lower arrow indicates unbound DNA. Quantitative PCR was performed for the positive control (PC) and negative control (NC) using DNA extracted from the gel and average cycle threshold values for each reaction are indicated below the corresponding bands. The `DESeq2` results for the FLANK library **C:** 1:0.2 and **D:** 1:0.1 *versus* unbound. The `DESeq2` results for the **ALL** library **E:** 1:0.2 and **F:** 1:0.1 *versus* unbound.

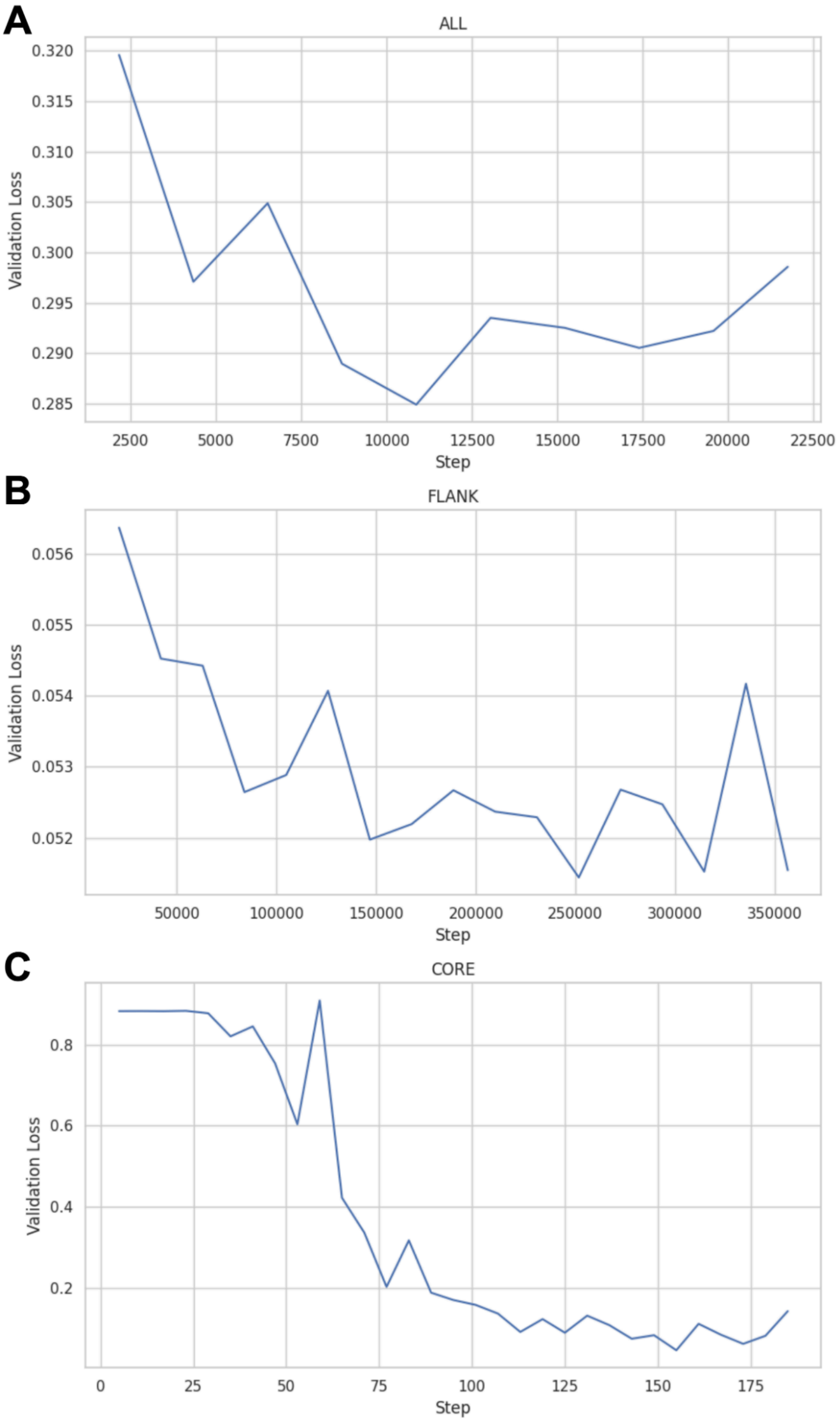


**Supplementary Figure S2:** The validation loss (mean squared error) for the VCNNBPNet models trained using three different libraries: **A: ALL**, **B: FLANK**, and **C: CORE**. The *x*-axis represents the training "Step" (indicating individual weight updates), while the *y*-axis shows the models' error as determined by the validation dataset.

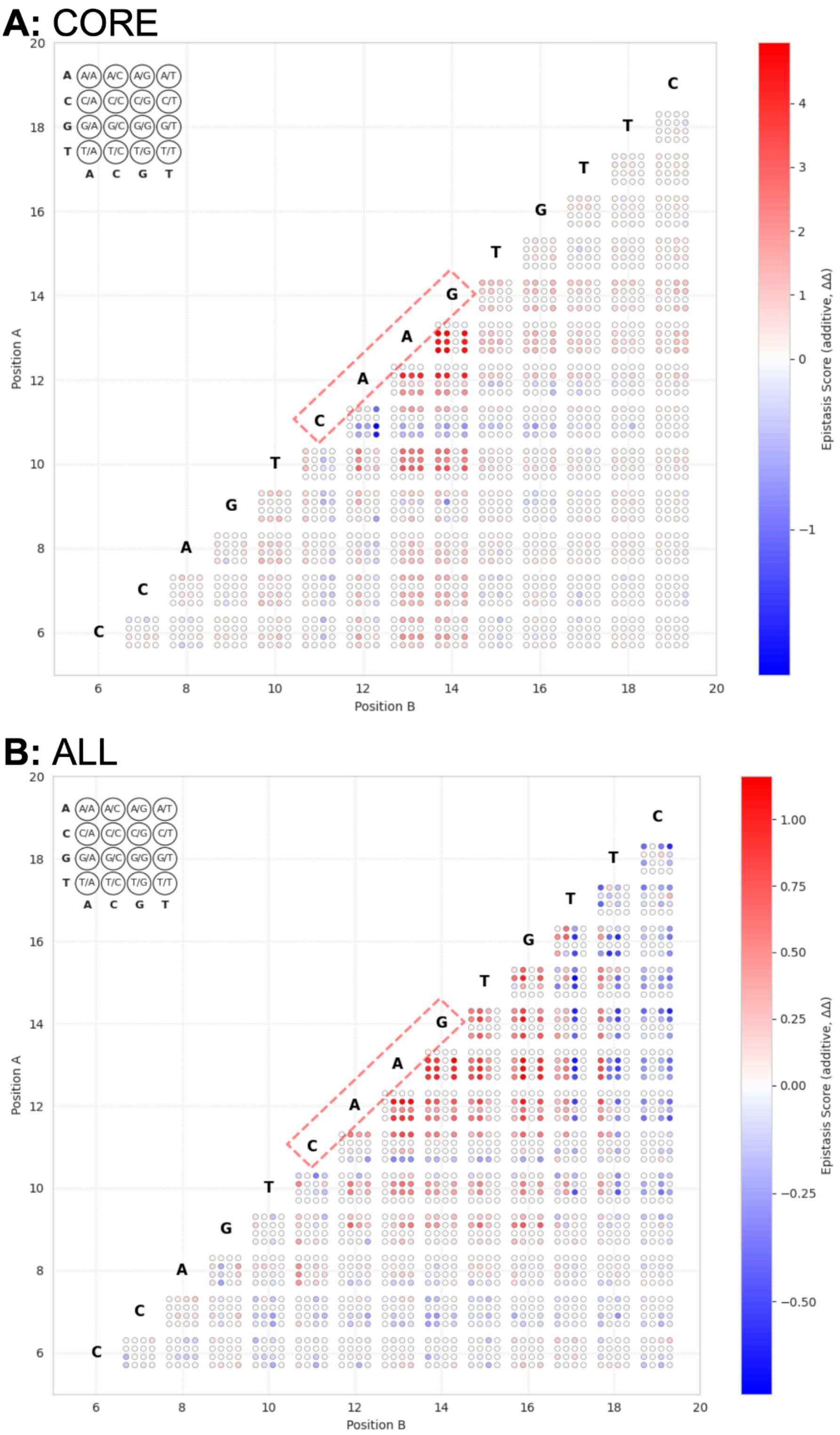


**Supplementary Figure S3**: Nucleotide interdependency map of the NKX2.1 binding site for the **A: CORE** and **B: ALL** models. The wildtype thyroglobulin promoter binding site sequence is shown diagonally with a red-dashed box enclosing the core nucleotides. For each double SNV, position A is on the y-axis and position B on the x-axis. The epistasis score is shown with a red-blue colour scale where a synergistic effect is red and an antagonistic effect is blue. The right side of the figure provides an example of how to read the map. Each square of 16 dots represents the variable combination of two nucleotide positions. Each possible nucleotide combination is depicted.
